# Transcriptional regulation of *HSFA7* and post-transcriptional modulation of *HSFB4a* by miRNA4200 govern general and varietal thermotolerance in tomato

**DOI:** 10.1101/2021.02.26.433069

**Authors:** Sombir Rao, Sonia Balyan, Jaishri Rubina Das, Radhika Verma, Saloni Mathur

## Abstract

Heat shock factors (HSFs) are at the core of heat stress (HS) response in plants. However, the contribution of HSFs governing the inherent thermotolerance mechanism in tomato from sub-tropical hot climates is poorly understood. With the above aim, comparative expression profiles of the *HSF* family in a HS tolerant (CLN1621L) and a sensitive cultivar (CA4) of tomato under HS revealed cultivar-biased regulation of an activator (HSFA7a) and repressor (HSFB4a) class HSF. Functional characterization of *HSFA7a* that was strongly up-regulated in the tolerant cultivar by VIGS-based silencing and overexpression established it as a positive regulator of HS-tolerance. While knock-down and overexpression analyses of *HSFB4a* that was down-regulated in CLN1621L in HS, showed it as a negative regulator of thermotolerance. Promoter:*GUS* reporter assays and promoter sequence analyses suggest heat-mediated transcriptional control of both the *HSF* genes in the contrasting cultivars. Moreover, we show *HSFB4a* is also regulated post-transcriptionally by microRNA Sly-miR4200 using degradome, short-tandem-target-mimic of Sly-miR4200 and transient *in-planta* Sly-miR4200-effector:*HSFB4a*-reporter assays. This miRNA is induced several folds upon HS in the tolerant variety thereby reducing *HSFB4a* levels. We thus propose that the alleviation of HSFB4a repressor governs thermotolerance in the tolerant cultivar by regulating downstream heat stress responsive genes.

## Introduction

The heat stress transcription factor (*HSF)* gene family members are the most critical regulators of transcriptional reprogramming during heat stress (Schramm et al. 2006, 2008; Ohama et al. 2016, Huang et al. 2016; Zang et al. 2019). Roles of HSFs have also been documented in response to other abiotic environmental cues (Nishizawa et al. 2006; Banti et al. 2010; Nishizawa et al. 2011; Bechtold et al. 2013; Liu et al. 2013; Tang et al. 2019; Zang et al. 2019), in plant development (Kotak et al. 2007; Waleed et al. 2018) and plant immune response to pathogens (Kumar et al. 2009; Bechtold et al. 2013; Wei et al. 2018). HSFs form the initial heat stress response (HSR) defense mechanism which mediate the induction of several heat stress responsive genes including heat shock proteins (HSPs) and HSF themselves by recognizing the heat stress elements (HSEs: 5’AGAAnnTTCT3’) in the promoter regions (Mishra et al. 2002; Schramm et al. 2008; Yoshida et al. 2008, 2011; Chen et al. 2010; Sato et al. 2014). This evolutionary conserved gene family is categorised into three classes (class A, B and C) based on its structural features (Scharf et al. 2012). The *HSF* family has expanded considerably during evolution in plants; there are 21 HSFs in Arabidopsis, 25 in rice, 27 in tomato and 52 in soybean (Scharf et al. 2012). Despite many conserved features, members of the *HSF* family show strong diversification of expression patterns and functions within the *HSF* family (von Koskull-Döring et al. 2007; Scharf et al. 2012; Xin et al. 2019). Along with Arabidopsis, tomato has also been used as a model plant to understand many aspects of HSF-mediated heat stress regulation (Rivero et al. 2001; Sato et al. 2006; Li et al. 2012, Hu et al. 2020a, 2020b).

Exposure of plants to heat stress leads to activation of master regulator *HSFA1s*, which trigger a transcriptional cascade composed of various *HSFAs* (*HSFA1e*, *HSFA2, HSFA3, HSFA7a*, and *HSFA7b*), *HSFBs* (*HSFB1*, *HSFB2a*, and *HSFB2b*), *DREB2A* and *MBF1c* (Mishra et al. 2002; Liu et al. 2011; Yoshida et al. 2011; Liu and Charng, 2013). These *HSFA1* induced HSFs include both enhancers as well as repressors of the HSR. Class A HSFs, *DREB2A* and *MBF1c* have been established as positive regulators of HSR (Sakuma et al. 2006; Larkindale and Vierling, 2008; Schramm et al. 2008; Yoshida et al. 2008; Suzuki et al. 2011; Nishizawa-Yokoi et al. 2011). Studies have established *HSFA2* as the most heat-inducible HSF (Busch et al. 2005) that has pivotal roles in the late phase of HSR as it maintains acquired thermotolerance (Schramm et al. 2006; Charng et al. 2007; Wunderlich et al. 2007). *HSFA2* is also shown to be involved in providing fitness to plants under other environmental stresses (Nishizawa et al. 2006; Banti et al. 2010). Using *HSFA2* knock-down tomato plants, Fragkostefanakis et al. (2016) have shown the role of *HSFA2* in regulating protective mechanisms in anthers during heat stress. Hu et al. (2020b) have illustrated the role of pre-mRNA splicing of *HSFA2* in governing thermotolerance in cultivated varieties during domestication. Strong induction in the transcripts of *HSFA6b* and *HSFA7* was observed in anthers of *HSFA2* knock-down tomato plants, Fragkostefanakis et al. (2016). *HSFA7* also exhibits strong up-regulation upon heat stress and plays key roles in cytosolic protein response (Busch et al. 2005; Charng et al. 2006; Cortijo et al. 2017; Sugio et al. 2009) and heat acclimation (Larkindale and Vierling, 2008). *HSFA6b* is involved in ABA-mediated salt and drought resistance and thermotolerance acquisition (Huang et al. 2016).

Heat tolerant and sensitive tomato varieties/genotypes have also been used to understand the naturally evolved signalling mechanisms of acquiring HS-resilience (Frank et al. 2009; Bita et al. 2011; Balyan et al. 2020; Hu et al. 2020a). These studies have provided a repertoire of genes involved in core and genotype-dependent HSR mechanisms in tomato. While Bita et al. (2011) used cDNA-AFLP coupled with microarray analyses in meiotic anther under moderate heat stress, Hu et al. (2020a) used seedlings from four genotypes exhibiting different thermotolerance and Balyan et al. (2020) performed RNA-seq in leaf from contrasting cultivar pair growing in the Indian sub-continent. Apart from canonical HSR regulators, several novel regulators of heat tolerance like Solyc09g014280 (*Acylsugar acyltransferase*), Solyc07g056570 (*Notabilis*) and Solyc03g020030 (*Pin-II proteinase inhibitor*) were functionally shown to have roles in regulating the variable degree of thermotolerance in contrasting tomato cultivars, the tolerant CLN1621L (henceforth called as CLN) and CA4 (sensitive) (Balyan et al. 2020). While all studies have been done utilising tomato genotypes from the temperate climate, in this study we uncover the involvement of HSFs in governing the heat tolerance and sensitivity using CLN and CA4 that grow in the sub-tropical climatic zones. qRT-PCR based expression profiling of HSFs in contrasting cultivars revealed the heat stressed induced regulation of fifteen HSFs in a cultivar biased manner. Furthermore, *HSFA7* was induced several folds higher in the tolerant variety than the sensitive one and *HSFB4a* was highly down-regulated in the tolerant cultivar. Functional studies utilizing VIGS and transient overexpression of these two HSF genes established *HSFA7* as positive regulator of HS and *HSFB4a* as a negative regulator of thermotolerance in tomato. Furthermore, we find cultivar biased heat mediated regulation of *HSFB4a* at both transcriptional and post-transcriptional levels. We show that *HSFB4a* is a target of microRNA, Sly-miR4200 that itself is regulated differentially between the cultivars. These results provide insights in to the unique and complex regulatory system of the HSR in plants.

## Materials and methods

### Plant material and stress conditions

Three-days-old seedlings of tomato (Solanum lycopersicum) cultivar, CLN1621L (CLN) and CA4 showing growth uniformity were transferred to soilrite filled plastic pots and placed in a plant growth chamber (CMP6050, Conviron, Canada) maintained at 60% relative humidity, 26 °C/21 °C (day/night: 16/8 h), light intensity 300 μM per m2 per sec. For heat stress 30 days old plants were initially exposed for 4 h at a gradual increasing range of temperature from 26 °C to 45 °C then exposed additionally for 4.5 h at 45 °C. Drought, salt and cold stress were applied as discussed by Paul et al. (2016). Leaves from these experimental plants were harvested, frozen in liquid nitrogen and kept at −80 °C until use.

### Real-time expression analysis

Total RNA was isolated from leaf tissues using Trizol (Ambion, USA) as per the manufacturer’s protocol. Total RNA was treated with DNase I (Ambion, USA) to remove genomic DNA contamination. For real-time PCR (qRT-PCR) analysis, 2 μg of total RNA was used for cDNA synthesis by using high-capacity cDNA reverse transcription kit (Applied Biosystem, USA) following the manufacturer’s protocol. The cDNA reaction mixture was diluted twice, and 0.5 μl was used as a template in a 10 μl PCR reaction. All the reactions were performed in a CFX-Connect real-time PCR detection system (Bio–Rad, www.bio-rad.com/) using GO Taq qPCR master mix (Promega, USA). Each experiment was repeated at least three times. The relative fold change was calculated by following the ddCT method using actin as normalization reference control (Qiu et al. 2016; Han et al. 2020; Yan et al. 2020). The primers used in this study are listed in Supplementary Table S1.

### Promoter sequence analysis

The gene and promoter (2kb) sequences of HSFB4a and HSFA7 were extracted from the genomes of CLN and CA4 (in house unpublished sequence data). The 2 kb sequences upstream of the translational start codon (ATG or CDS start site) of tomato HSFs were also downloaded from SGN. The pair-wise sequence alignments were performed using Alignment and trees tool of CLC genomics workbench (Qiagen). The promoter sequences were analyzed for the presence of putative cis-regulatory elements by PlantCARE (Rombauts et al. 1999) and/or PlantPAN 3.0 (Chow et al. 2019). To identify the probable TFs having TFBS in the promoter, the protein sequence of PLANTPAN identified Arabidopsis TFs were used as query against the S_lycopersicum_protein _database version 4 using BLASTP and the topmost hit was considered as the orthologue TF. Further the expressions of above candidates were extracted from the published transcriptome data of CLN and CA4 (Balyan et al.2020).

### Thermotolerance and growth measurement assay for tomato seedlings overexpressing HSFs

The transient overexpression of HSFA7 and HSFB4a genes followed by thermotolerance assays were standardized for different tomato cultivars by following the methodology adopted by Queitsch et al. (2000) with modifications as discussed by Balyan et al.2020. Experiment was repeated four times with similar parameters with sample size of 70 seedlings per replicate.

### Tobacco Rattle Virus-induced gene silencing (VIGS)

Transient silencing assays following the virus induced gene silencing approach (VIGS) was performed as described by Senthil and Mysore (2014). For the silencing of HSFA7 and HSFB4a genes, 300 to 400 bp cDNA fragment selected by the VIGS tool (Sol Genomics Network) was amplified and cloned into TRV2 vector. After confirmation by Sanger sequencing, the TRV2-gene vectors were transformed into Agrobacterium tumefaciens strain GV3101. About 15 days-old tomato plants were inoculated as described previously (Senthil and Mysore, 2014). TRV2 Empty vector infiltrated plants were used as a control. TRV-EV and TRV-HSFs inoculated plants were placed in moist and dark conditions for 3 days and then transferred to light; after 3 weeks, infected plants were subjected to HS at 45°C for 4.5h. Establishment of VIGS was confirmed by detecting viral movement protein in TRV-silenced plants by PCR using genomic DNA as template. TRV-based tomato Phytoene desaturase (PDS) silencing was used to confirm establishment of VIGS-mediated silencing. Both the TRV-PDS and TRV-HSFs batches were started and inoculated at the same time. Appearance of white patches on leaves of TRV-PDS silenced plants confirms the onset of silencing in both the batches. Furthermore, silencing of HSFs was also confirmed by quantifying its transcript levels in silenced plants as compared to empty TRV vector containing plants. Leaves were collected and frozen for qRT-PCR analysis. Survival and cell death were assayed after 6 days recovery post HS. The experiment was performed in three biological replicates with 8 plants per replicate for each gene.

### Measurement of gas-exchange parameters

Gas-exchange measurements, including net photosynthetic rate (A), water use efficiency (WUEi), transpiration rate (E) and stomatal conductance (Gs) were measured simultaneously by using a portable LICOR 6400 photosynthesis system (LI-6400, Li-Cor Inc., Lincoln NE, USA) as discussed by Balyan et al.2020.

### Histochemical detection of H2O2 and cell death assay

Histochemical staining was done by using 3,3’-diaminobenzidine (DAB) stain to detect hydrogen peroxide (H2O2) (Daudi et al. 2012) and cell death was measured by using trypan blue as described previously (Koch and Slusarenko, 1990). For these assays 6th and 7th leaf from 6-week-old pTRV and pTRV-HSFs plants were stained immediately after HS (45°C for 4.5h). Experiments were repeated three times with similar parameters with sample size of 8 plants per replicate.

### GUS aided promoter reporter analysis of HSFs

For Pro-HSFA7:GUS and Pro-HSFB4a:GUS, 2000 bp region upstream of the translation start codon was amplified from genomic DNA and cloned into pbi101 vector and transformed in A. tumefaciens EHA105 strain. Tomato seedlings were transformed as per the above-mentioned protocol for transient assays. After 2 days of infiltration transformed seedlings were subjected to different stress treatments. For heat stress treatment, transformed seedlings were kept at 45°C for 2h. Seedlings were subjected to GUS staining post stress treatments and analyzed for GUS expression by qRT-PCR as per Rao et al. 2020.

### Prediction of HSFB4a as miRNA target by degradome analysis

The prediction of miRNAs targeting HSFB4a was done by analysing the PARE datasets (GSM1047560, GSM1047562, GSM1047563, GSM553688, GSM553689, GSM553690, GSM1379695, GSM1379694, GSM1372427 and GSM1372426) of different tissues of tomato from NCBI (https://www.ncbi.nlm.nih.gov/sra). Briefly, all the PARE datasets was trimmed using Trim adapter tool of CLC Genomics workbench version 9. All the mature miRNAs sequences of tomato were used as miRNA query against the tomato gene model sequences of tomato (ITAG version 4) following the CleaveLand4 pipeline (Addo-Quaye et al. 2009) with the default settings. Further the miRNA/mRNA pairs with an alignment score ≤ 7, total PARE reads more ≥2 being cut at 10th position (+/− 1 base) with evidence from at least 2 datasets were considered as targets.

### Agro-infiltration and transient assay for confirming HSFB4a as miRNA target

Agro-infiltration was performed in 4-week-old Nicotiana benthamiana leaves for target validation. Sly-MIR4200-OE and GFP-HSFB4a overexpression constructs were transformed into Agrobacterium tumefaciens EHA105 strain. For infiltrations, overnight cultures of individual constructs were harvested and suspended at OD 1 in infiltration buffer (10 mM MgCl2, 10 mM MES, pH 5.8, 0.5% glucose and 150 μM acetosyringone) and incubated for 3 h at room temperature. A set of N. benthamiana leaves were infiltrated by syringe with the target constructs alone or in a 1:1 ratio of miRNA precursor-overexpression:target (effector:reporter) constructs. HPT-II gene in vector was used to normalize target abundance in qPCR experiments.

### Short-tandem-target-mimic (STTM) to sponge miR-4200 in tomato

STTM-4200 suppression construct was generated as discussed in Rao et al. (2020). The construct was transformed into Agrobacterium tumefaciens strain EHA105, and vacuum infiltrated in 5 days’ tomato seedlings. miR4200 chelation was confirmed by measuring HSFB4a transcripts levels by qRT-PCR.

## Results

### Distinct modulation of heat stress transcription factors in tolerant (CLN) and sensitive (CA4) tomato cultivars during heat stress

A comparative transcriptome analysis during heat stress from our laboratory has previously noted an inherent cultivar-specific orchestration of 402 transcription factors belonging to 46 families in regulating heat responsive genes in the heat tolerant (CLN) and the heat sensitive (CA4) tomato cultivars (Balyan et al. 2020). RNA-seq analysis of the master-regulator HSF family members during heat stress showed similar to differential accumulation of *HSF* transcripts in contrasting tomato cultivars suggesting the possible involvement of HSFs in governing varietal heat tolerance and sensitivity (Balyan et al. 2020). Taking cues from this observation, we extended the study by analyzing the expression of the 26 tomato HSFs (Scharf et al. 2012) in the tolerant and the sensitive cultivars by qRT-PCR during HS. We find that fifteen tomato *HSF* genes exhibit HS-mediated up-regulation in both the cultivars (Figure 1a, b and Supplementary table S2). The highest elevation is seen for *HSFA2,* it being nearly 157 fold and 90 fold up-regulated in CLN and CA4, respectively (Figure 1a, Supplementary table S2). *HSFA4b* is down-regulated under HS in CLN but up-regulated in CA4 (Figure 1a). The expression of two HSFs (*HSFA1a* and *HSFB3a*) remains unchanged in both the cultivars upon heat stress. Five HSFs (*HSFA1c, HSFA1e, HSFA4a, HSFA4b, HSFA6a*) exhibit significant up-regulation in sensitive cultivar CA4. Among these five HSFs up-regulated in CA4, four HSFs remain non-differential in CLN while *HSFA4b* shows significant down-regulation (Figure 1a). Moreover, we find that the activator class HSF (*HSFA7*) exhibits strong up-regulation (~30 fold), while the repressor class B HSF (*HSFB4a*) shows strong down-regulation in the tolerant cultivar, CLN. On the other hand, *HSFA7* is only 4-fold up-regulated and the transcript abundance of *HSFB4a* remains unchanged in response to HS in the sensitive cultivar (CA4) (Figure 1a). Based on this observation, we selected *HSFA7* and *HSFB4a* for further characterization in governing heat stress response.

**Figure 1:**
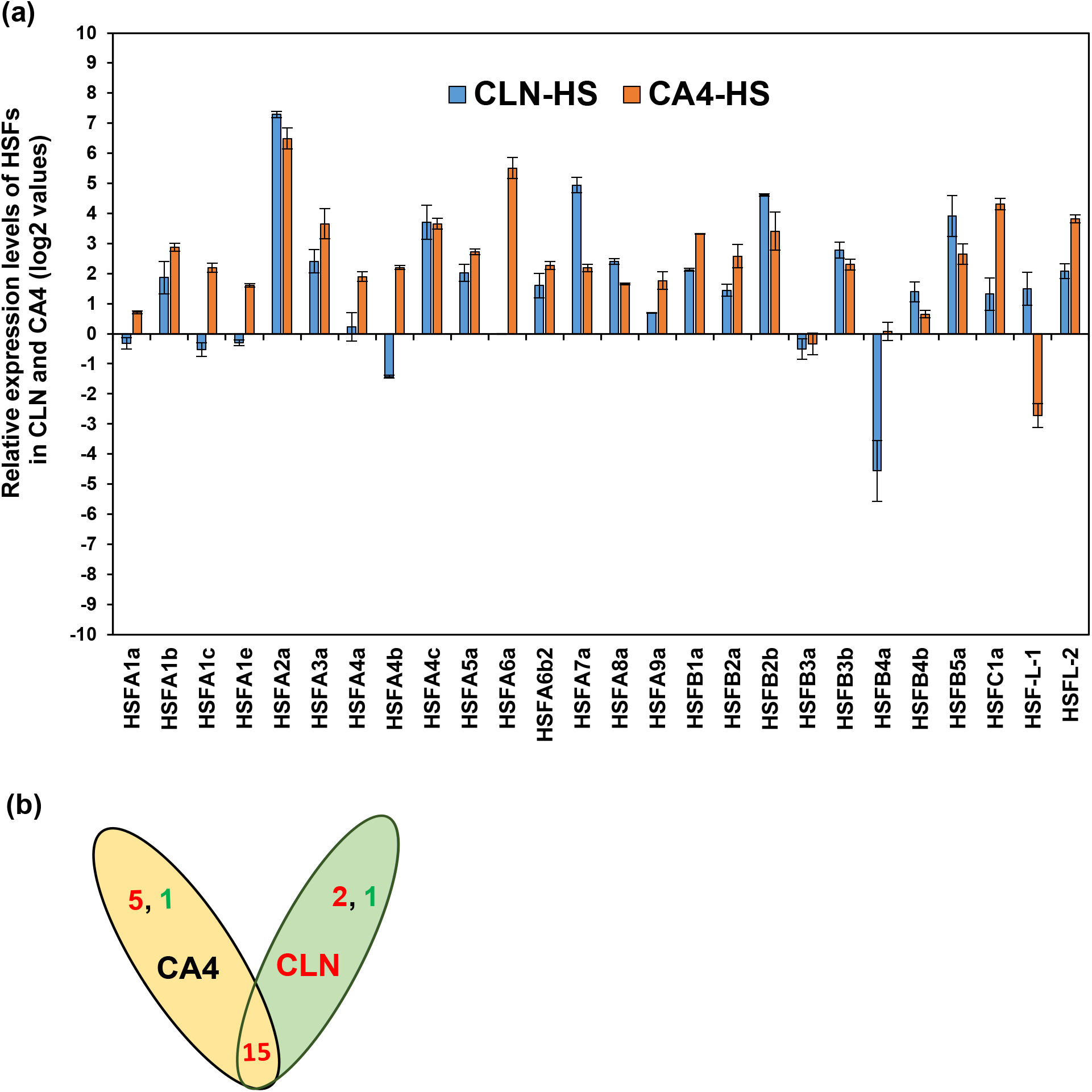
Expression profiling of HSFs in leaf of contrasting tomato cultivars upon heat stress. (a) qRT-PCR based expression analysis of HSFs during HS in leaf of CLN (tolerant) and CA4 (sensitive) cultivar. *Actin* was used as the normalization control. Fold change of *HSFs* during heat stress was calculated by setting the fold change value of CLN and CA4 plants kept in control conditions as one. Log2 transformation was applied to the fold-change data to obtain negative fold change values. Experiment was repeated three times and average values are plotted as bars, error bars depict standard error between three replicates. (b) Venn diagram depicting shared and uniquely upregulated and downregulated HSFs in contrasting cultivar pair during HS. Red coloured numerals depict the number of genes that are up-regulated, while green coloured numerals depict down-regulated HSF genes.

### *HSFA7* is critical for high temperature tolerance in tomato

Real-time based expression analysis of HSFs in CLN and CA4 (Figure 1) revealed the strong induction of two Class A *HSF* genes viz., *HSFA2* and *HSFA7* in heat stress. These HSFs show higher degree of induction in tolerant cultivar CLN as compared to sensitive cultivar CA4 (Figure 1a), this induction is nearly 7-fold more in CLN than CA4 for *HSFA7*. *HSFA2* has been extensively studied and established as a key player of thermotolerance at both vegetative and reproductive stages (Giorno et al. 2010; Fragkostefanakis et al. 2016). On the other hand, the activator and regulatory functions of *HSFA7* during heat stress response is still limited. We thus, explored the possible roles of *HSFA7* in regulating heat stress tolerance of tomato. qRT-PCR based comparative expression profiling of HSFs in non-stressed CA4 and CLN plants revealed the presence similar inherent levels of HSFA7 transcripts in both the cultivars (Supplementary figure S1). Thus, imposition of heat stress orchestrates the transcriptional enhancement of HSFA7 in both the cultivars (Figure 1a), suggesting the involvement of HSFA7 in general thermotolerance pathway. To further confirm the heat-mediated induction of *HSFA7* transcription *in-planta,* we performed *GUS* aided promoter reporter assay of *HSFA7* by transforming *pHSFA7:GUS* sensor construct in tomato (Supplementary figure S2a, b). Enhanced *GUS* expression in HS upon histochemical staining (Supplementary figure S2a) as well as increased accumulation of *GUS* transcripts in *pHSFA7:GUS* transformed seedlings in response to HS (Supplementary figure S2b), confirmed the heat-mediated induction of *HSFA7* transcription.

Furthermore, to explore whether the heat induced differential upregulation of *HSFA7* in CLN-CA4 cultivars is because of the sequence variation in promoter of the two cultivars. Comparative analysis of promoter sequences for both the varieties highlighted no SNPs (Supplementary figure S3), thus a *cis*-sites mediated regulatory influence in the contrasting expression was ruled out and pointed towards a probable TF-mediated cultivar-specific regulation. Taken together, the qRT-PCR based *HSFA7* expression data and the *pHSFA7*:*GUS* assays revealed the strong up-regulation of *HSFA7* (~30-fold) during heat stress in heat tolerant cultivar CLN (Figure 1a, Supplementary figure S2), suggesting its crucial role in governing heat tolerance. Therefore, to further explore the functions of *HSFA7* in governing heat tolerance, virus induced gene silencing (VIGS) approach was utilized to knock-down the expression of *HSFA7* in tomato plants. Thermotolerance of TRV-*HSFA7* silenced plants was assessed after 3-weeks of agro-infiltration and the establishment of gene silencing was confirmed by strong reduction in the *HSFA7* mRNA levels (~64%) and appearance of bleaching phenotype in a parallel set of *phytoene desaturase* (*PDS)* silenced plants, used as VIGS silencing control (Supplementary figure S4a, b). We have previously shown the specificity of *HSFA7* down-regulation in TRV-*HSFA7* silenced plants, by examining the expression profiles of putative off targets as discussed by Rao et al. (2021). For thermotolerance assay, TRV-*HSFA7* silenced plants were given HS and scored for survival six-days-after HS imposition. Higher degree of damage was observed in leaves (burning and drooping) of TRV-*HSFA7* silenced plants than the vector control plants (Figure 2a). The TRV-*HSFA7* knock-down plants showed nearly 6-fold higher mortality rate than vector control plants as only ~5.6% TRV-*HSFA7* knock-down survived (judged by phenotype of the 7th to 9th leaves and emergence of new leaves) in comparison to ~36.6% for vector-control plants (Figure 2b). These results suggest that *HSFA7* positively regulates thermotolerance in tomato. Survival and growth measurement assays of seedlings overexpressing *HSFA7,* under HS further strengthened our findings about the involvement of *HSFA7* in positively regulating thermotolerance in tomato. The P35S:*HSFA7* over-expressing seedlings are more robust and healthier phenotypically than the vector control plants post heat stress (Figure 2c). The seedling survival increases by nearly 2-fold from 40% in vector control to ~80% in P35S:*HSFA7* over-expressing plants (Figure 2d), the hypocotyl length under HS also doubles in over-expression plants as compared to vector control plants (Figure 2e, f). In addition, seedlings over-expressing P35S:*HSFA7* also assimilate higher biomass in comparison to vector control plants during heat stress (Figure 2g), reiterating the positive role of *HSFA7* in governing thermotolerance.

**Figure 2:**
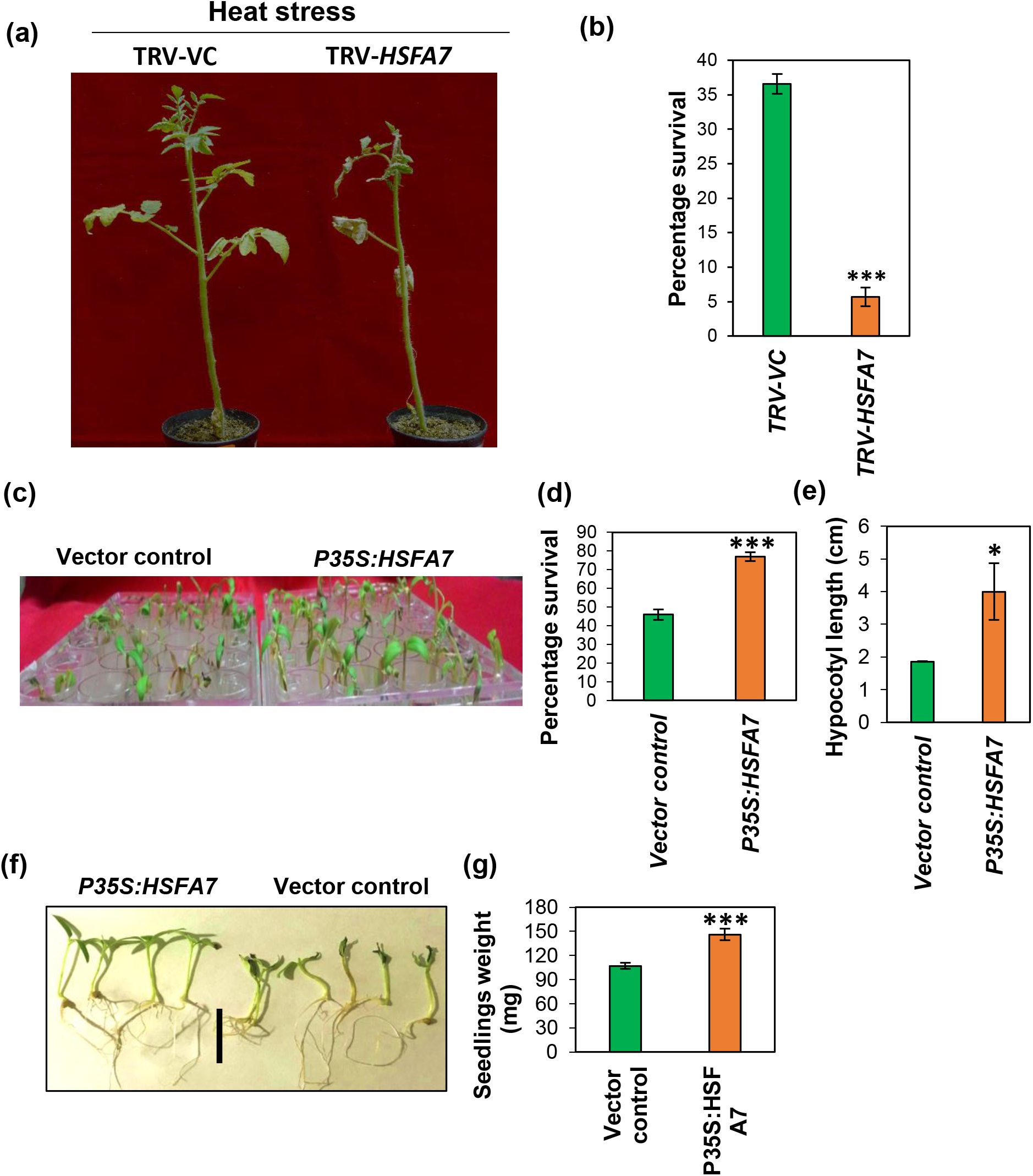
*In-planta* validation of HSFA7 functions in heat stress tolerance of tomato. (a) TRV-VC and *TRV-HSFA7* plant phenotypes after heat stress (HS) treatment. (b) Percentage survival of TRV-VC and *TRV-HSFA7* plants after heat stress. (c) Phenotype of *P35S:HSFA7* overexpressing and empty vector control seedlings after heat stress. (d) Estimation of percentage survival and (f) hypocotyl length after HS in *P35S:HSFA7* overexpressing and empty vector control seedlings. (g) Estimation of seedling weight in empty vector and *TRV-HSFA7* VIGS silenced plants after heat stress. Data are means and standard error of four biological sets of 70 seedlings each. *p<0.05 and ***p<0.001

### Differential regulation of *HSFB4a* transcription in CLN and CA4 under control and heat stress

Class B HSFs are known as transcriptional repressors that negatively regulate thermotolerance, as shown in the higher acquired thermotolerance property of the *hsfb1-hsfb2b* double knockout Arabidopsis mutant plants (Kumar et al. 2009; Ikeda et al. 2011). This study finds that *HSFB4a* expression is highly reduced in CLN during control as well as in HS, while its expression remains unchanged in CA4 (Figure 1a, Supplementary figure S1). To further delineate the underlying mechanism for this differential accumulation of *HSFB4a* transcripts between the two varieties, we compared their promoter sequences (2 kb upstream region of ATG). The *HSFB4a* promoter sequences of both the cultivars is identical (Supplementary figure S5), thus the *cis*-sites mediated regulatory influence in the promoters for the observed differential expression levels was ruled out. However, when we measured the transcripts levels of *GUS* gene hooked to the 2 kb CLN and CA4 *HSFB4a* promoters in CLN and CA4 background respectively, we find stark difference in their transcriptional potential under heat stress. The promoter reporter assay highlights that *GUS* expression is up-regulated by ~2.5 fold in pCA4-*HSFB4a* in CA4 background while there is strong reduction of reporter gene expression (~9 folds) in CLN seedlings infiltrated with pCLN-*HSFB4a* (Supplementary figure S6). This differential transcriptional regulation is also evident from the *GUS* histochemical assay in seedlings upon heat stress (Supplementary figure S6b). This analysis confirms the existence of regulation of *HSFB4a* at transcription level during HS in a cultivar-biased manner. These results point towards a probable TF-mediated differential regulation between the two varieties. To search for the TFs regulating the above observed differential transcriptional regulation of *HSFB4a*, the promoter was analysed using PLANT-PAN database that showed 46 potential TF binding sites with 69 candidate TFs (Supplementary table S3a). Using RNA-seq based differential analysis of TFs in response to HS (Balyan et al, 2020), 27 of the 69 TFs are HS responsive (Supplementary table S3b). However, only three TFs (Solyc03g006830, MADS box; Solyc08g076930, jasmonic acid 3 and Solyc01g095030, Altered Response to Salt Stress 1) were significantly altered in CLN and non-differential in CA4 (Supplementary table S3b). Whether this cultivar-based TF difference accounts for the differential activity of *HSFB4a* needs to be ascertained.

### *HSFB4a* is post-transcriptionally regulated by Sly-miR4200

The qRT-PCR based expression analysis during HS reveals strong down-regulation (~ - 23-folds) of *HSFB4a* in CLN (Figure 1a), while *in-planta GUS* aided promoter reporter assay of pCLN-*HSFB4a* during HS in CLN background exhibit only 9-fold down-regulation (Supplementary figure S6a, b). Similarly, the expression profiles of *HSFB4a* in CA4 exhibit non-differential expression in response to heat (Figure 1a), while *in-planta* promoter:*GUS* reporter assay of pCA4-HSFB4a during HS in CA4 background exhibits significant up-regulation (~2.5 folds) (Supplementary figure S6a, b). To address this apparent discrepancy in the results obtained by qRT-PCR and promoter reporter assays, we examined the involvement of post-transcriptional regulation on this HSF by microRNAs. A search for possible miRNAs targeting *HSFB4a* was done via degradome analysis by running CleaveLand (version 4) pipeline on 10 publically available PARE (parallel analysis of RNA ends) libraries of tomato downloaded from NCBI. Our search yielded Sly-miR4200 as a putative regulator of *HSFB4a* post-transcriptionally (Figure 3a) by *HSFB4a* transcript cleavage (Figure 3a, b). It has been highlighted by many authors that many spurious miRNAs have been classified in literature. We therefore, checked the precursor structure of *Sly*-*MIR4200* using online RNA fold tools. *Sly*-*MIR4200* showed a valid stem-loop structure that fulfilled all the defined guidelines for it to be a bonafide precursor (Figure 3b). Furthermore, to confirm and validate the miRNA-mediated cleavage of *HSFB4a in-planta*, *Nicotiana benthamiana* based agro-infiltration transient assay was performed. The putative target (*HSFB4a)* was cloned in frame with *GFP* and expressed under CaMV35S constitutive promoter (target:reporter construct) in leaves with or without *Sly-MIR4200* precursor effector plasmid. *GFP* levels were quantified and normalized with *hygromycin* (*HPT-II*) gene that is co-expressed from the vector. Reduction in the *GFP* levels (~2.5 fold) when the effector plasmid is co-infiltrated in leaves confirms *HSFB4a* as Sly-miR4200 target (Figure 3c). To further confirm the miR-4200-mediated cleavage of *HSFB4a*, miR-4200 expression was suppressed transiently by sponging-up mature miRNA by short-tandem-target-mimic (STTM) expressed under 2 × 35S promoter in tomato. Transcript levels of *HSFB4a* in *STTM-MIR4200* infiltrated plants showed significant elevation confirming it as true miR-4200 target (Figure 3d).

**Figure 3:**
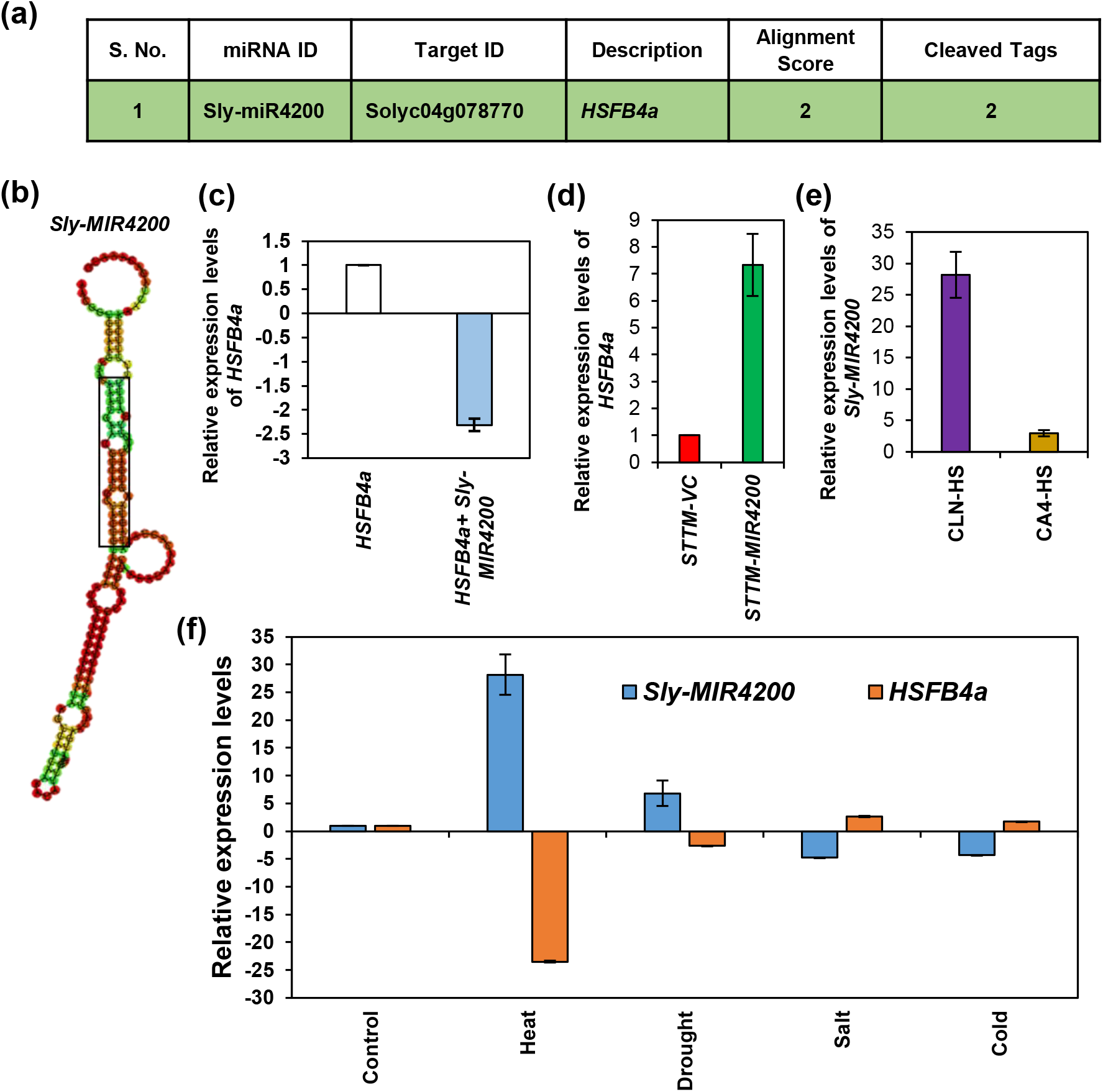
Post-transcriptional regulation of *HSFB4a* by Sly-miR4200 during heat stress. (a) *HSFB4a* identified as target of Sly-miR4200 by degradome analysis. (b) The secondary stem loop structure of Sly-miR4200 as predicted using the RNAfold web server. The mature miRNA/miRNA* portion is marked by a black box. (c) Validation of *HSFB4a* as Sly-miR4200 target by transiently overexpressing *HSFB4a* target (target-reporter) and Sly-miR4200 precursors (miRNA effector). The relative expression levels of targets were assessed by comparing the expression of the GFP fused to miRNA-sensitive target alone and when co-infiltrated with different miRNA in *Nicotiana benthamiana* leaves. qRT-PCR analysis of *GFP* mRNAs of p35S:GFP:targets was performed with *hygromycin* gene (*HPT-II)* that is present in the same vector for normalization. Data is shown as means ± Standard error of three biological replicates with 3-4 agro-infiltrated plants for each construct. (d) *In-planta* validation of *HSFB4a* as Sly-miR-4200 target by using STTM (Short-tandem-target mimic). Enhanced expression of *HSFB4a confirms* the miRNA chelation as measured by qRT-PCR. (e) Expression levels of *Sly-MIR4200* in CLN and CA4 during heat stress. (f) Expression profiles of *Sly-MIR4200:HSFB4a* module in different abiotic stresses in tomato. *Actin* was used as the normalization control. Experiment was repeated three times and average values are plotted as bars, error bars depict standard error between three replicates in Figure (d-f).

### *Sly-MIR4200*:*HSFB4a* module is orchestrated in a cultivar-specific manner during heat stress

Furthermore, we asked the question, whether *Sly-MIR4200* is also regulated in a cultivar-biased manner like *HSFB4a* transcripts. Expression analysis of *Sly-MIR4200* was carried using qRT-PCR in heat treated leaf of CLN and CA4 plants. The expression profiling revealed that *Sly-MIR4200* exhibits strong induction (at least 28-fold) during HS in CLN (Figure 3e), and also maintained at abundant level during control conditions in CLN leaf (Supplementary figure S7, Supplementary table S4). In contrast, *Sly-MIR4200* exhibits low abundance in CA4 leaf during control as well as HS conditions (Ct values above 35) and is ~2-fold enhanced (Supplementary figure S7, Supplementary table S4, Figure 3e). Thus, there is varietal-biased expression of *Sly-MIR4200* that in turn modulates the expression of *HSFB4a* in a cultivar-specific manner during HS. Notably, the CA4 *HSFB4a* promoter shows significant enhancement in transcriptional rate (Supplementary figure S6a, b) however, the quantification of *HSFB4a* transcripts during HS in CA4 by qRT-PCR exhibits no differential expression (Figure 1a). We hypothesize that the slight up-regulation of *Sly-MIR4200* (Figure 3e) maintains the *HSFB4a* transcripts levels post-transcriptionally during HS. In this way, the differential expression of Sly-miR4200 between CLN and CA4 contributes towards the maintenance of differential *HSFB4a* transcript levels in the heat tolerant and sensitive cultivar.

To further elucidate the existence of *Sly-MIR4200*:HSFB4a module in tomato leaf during different abiotic stresses, we carried out a qRT-PCR based comparative expression profiling of *Sly-MIR4200* precursor and *HSFB4a* transcripts. The *Sly-MIR4200* and *HSFB4a* transcripts exhibits inverse expression patterns during heat, drought, and salt stress (Figure 3f), suggesting the possible involvement of this module during abiotic stress responses. Of note, this *Sly-MIR4200:HSFB4a* module is most prominent during heat stress (Figure 3f).

### *HSFB4a* negatively regulates high temperature tolerance in tomato

To establish whether *HSFB4a* abundance has any direct correlation with CA4 heat stress sensitivity, we did functional characterization of *HSFB4a* utilizing VIGS to knock-down gene expression as well as transient over-expression of P35S:*HSFB4a* in tomato seedlings. *HSFB4a* was transiently over-expressed in 5-days-old tomato seedlings under CaMV35S constitutive promoter. Measurement of seedling growth by gauging hypocotyl length and dry weight during control and heat stressed conditions confirm *HSFB4a* as a repressor of growth. Tomato seedlings overexpressing *HSFB4a* exhibit significantly reduced hypocotyl length and dry weight as compared to vector control plants during non-stressed conditions (Figure 4a, c, d). Furthermore, thermotolerance assays of *HSFB4a* confirm *HSFB4a* as a repressor of heat tolerance as seedlings over-expressing *HSFB4a* (P35S:*HSFB4a*) exhibit poor growth, burnt cotyledons and reduced survival rate as compared to vector control plants 6-days-post heat stress treatment (Figure 4a-d). Survival rate of P35S:*HSFB4a* over-expressing seedlings was reduced from 58% to 47% as compared to vector control plants (Figure 4b). Seedlings over-expressing P35S:*HSFB4a* also exhibit reduced hypocotyl length (Figure 4c) and reduced biomass (dry weight) (Figure 4d) as compared to vector control plants after heat stress. Furthermore, to concrete our findings, we performed VIGS of *HSFB4a* in tomato. Plants exhibiting successful knock-down (~40-fold reduction) for *HSFB4a* transcripts (Figure 4e) were further subjected to HS at 45 °C for 4.5 h and survival percentage as well as HS-mediated phenotypic differences were scored 6-days-post HS. To assess the specificity of *HSFB4a* silencing and to rule out the off target mediated alteration in post HS phenotyping and survival scoring, we predicted the putative off targets for *HSFB4a* by using VIGS tool available at SGN. Since no significant off-targets were shown by the VIGS-tool, we selected *HSFB4b* which belongs to the same subclass as HSFB4a as a probable off-target. qRT-PCR based expression profiling of *HSFB4b* revealed the unaltered expression pattern in *HSFB4a* VIGS silenced plants, confirming the specificity of silencing (Supplementary figure S8). The phenotypic assessment revealed that plants with silenced *HSFB4a* exhibit no HS severity in terms of damage and drooping of leaves (Figure 4f) and have nearly double survival rate in comparison to vector control plants (Figure 4g). Furthermore, we measured various physiological and photosynthesis parameters of TRV-*HSFB4a* silenced plants. Li-COR 6400 portable photosynthesis measuring system based measurement of gas exchange parameters after 6 days of the heat treatment revealed that, TRV-*HSFB4a* silenced plants exhibit significantly enhanced water use efficiency (WUEi) and net photosynthesis rate as well as reduced transpiration rate in comparison to vector control plants (Figure 4h) highlighting the role of *HSFB4a* as a negative regulator of thermotolerance. HS is associated with reactive oxygen species (ROS) burst and cell death. Silencing of *HSFB4a* shows reduced cell death (Figure 4i) and exhibits low levels of ROS accumulation as compared to vector control plants (Figure 4j). Thus, it appears that HS-mediated alleviation of *HSFB4a* in tolerant cultivar (CLN) governs its heat tolerance, while its abundance in CA4 negatively regulates its tolerance and leads to heat sensitivity.

**Figure 4:**
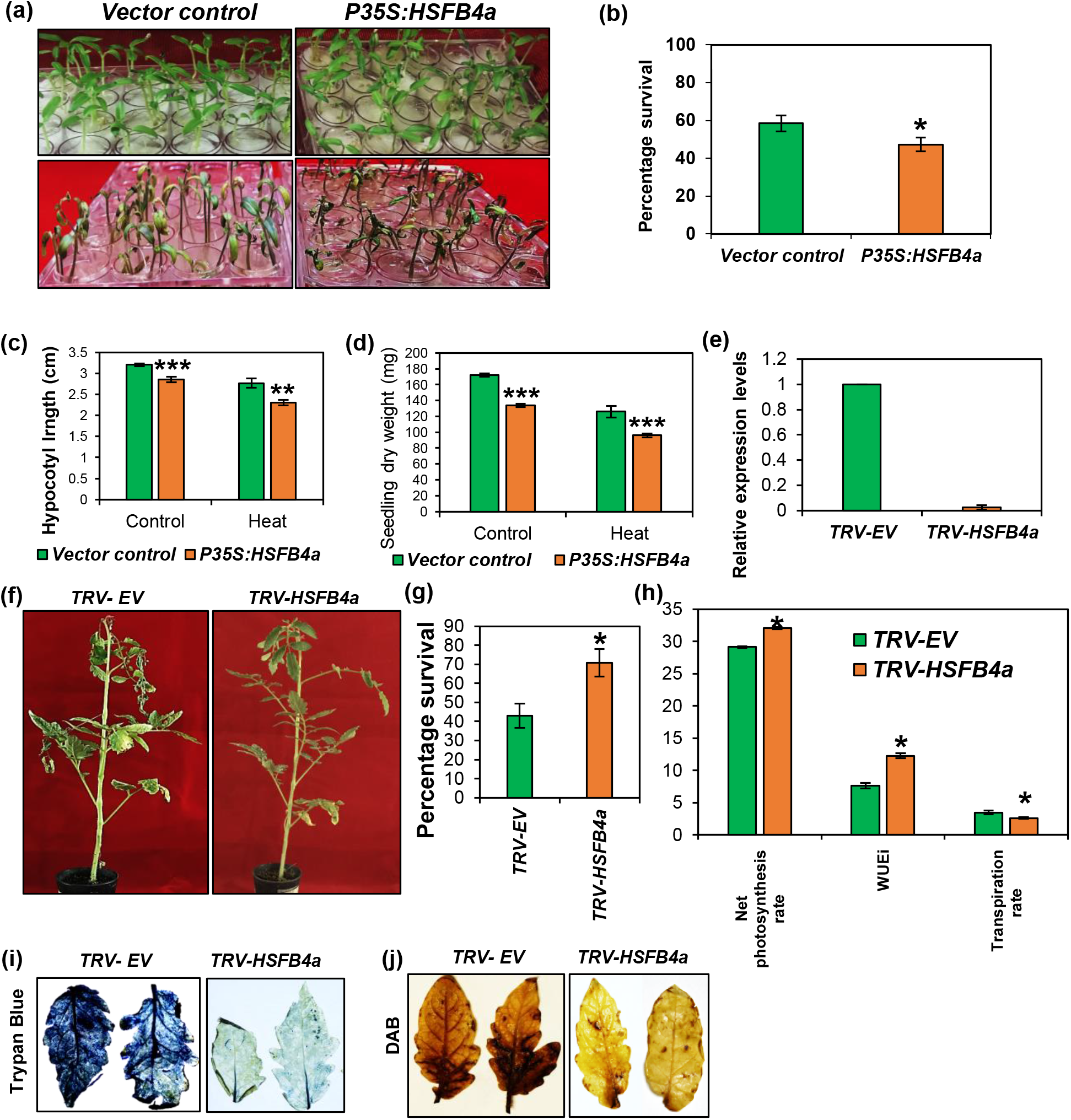
*In-planta* validation of *HSFB4a* as negative regulator of heat stress tolerance in tomato. (a) Phenotype of *P35S:HSFB4a* overexpressing and empty vector control seedlings in control and after heat stress (HS) conditions. (b) Estimation of percentage survival and (c) hypocotyl length in control and after HS in *P35S:HSFB4a* overexpressing and empty vector control seedlings. (d) Dry weight estimation in empty vector and *P35S:HSFB4a* overexpressing seedlings during control and after heat stress. Data are means and SE of four biological sets of 70 seedlings each. (e) Quantitative RT-PCR analysis of *HSFB4a* in TRV-EV and *TRV-HSFB4a* silenced plants confirming the silencing of *HSFB4a*. The expression levels of *HSFB4a* were calculated using the 2−ΔΔCt method and presented using fold-change values as compared to control. Fold change was calculated by setting the fold change values of *HSFB4a* in *TRV-EV* plants as one (f) TRV-EV and *TRV-HSFB4a* silenced plants phenotypes after heat stress treatment. (g) Percentage survival of HS treated TRV-EV and *TRV-HSFB4a* plants. (h) Measurement of net photosynthesis rate (μmol m−2s−1), water use efficiency (mmol mol−1) and transpiration rate (mmol m−2 s−1) in TRV-EVand *TRV-HSFB4a* silenced plants following heat stress. (i) Trypan blue and (j) DAB staining of HS treated leaves of TRV-EV and *TRV-HSFB4a* silenced plants. Plants were given heat stress 3 weeks post agro-infiltration. Survival was gauged 6 days post recovery. *p<0.05, **p<0.01 and ***p<0.001

### Heat tolerant CLN plants show enhanced heat-responsive gene expression

To further elucidate the molecular mechanism governing the thermal tolerance of CLN variety and to know whether the strong up-regulation of activator *HSFA7* and reduced expression of repressor *HSFB4a* during heat stress resulted in the higher accumulation of actual heat stress effectors proteins like HSPs and ROS scavenger enzyme like Ascorbate peroxidase (APX), qRT-PCR was performed. Expression profiling of heat stress responsive genes (HSPs and *APX*) in CLN and CA4 revealed that there is higher abundance of *HSP90, APX* and *HSP100* genes in CLN during control conditions (Figure 5a). Furthermore, upon imposition of heat stress, there is strong up-regulation of *sHSP17.6C-II, HSP90, APX* and *HSP70* in CLN as compared to the sensitive variety CA4 (Figure 5b). The heat-mediated up-regulation of HSR genes is also evident in CA4 but the expression levels are very strong and more pronounced in the tolerant cultivar (Figure 5b). The differential expression of the ROS scavenger *APX* during control and heat stresses CLN and CA4 prompted as to explore the levels of ROS production in the CLN and CA4 cultivars. DAB staining of CLN and CA4 during control conditions revealed the production of high levels of ROS in CA4 in comparison to the tolerant cultivar CLN (Figure 5c). The imposition of heat stress leads to the production of significant ROS in CLN cultivar, but its more evident in the sensitive cultivar CA4 (Figure 5d). Taking together the levels of APX and ROS, our analysis points toward the involvement of HSR genes in maintaining the tolerance of CLN during heat stress. Further, to relate the roles of strong transcriptional induction of *HSFA7* in CLN with its higher HSR expression levels, we checked the expression of HSR genes in *TRV-HSFA7* silenced plants. The qRT-PCR based expression profiling shows the loss of heat mediated regulation of HSR genes in *TRV-HSFA7* silenced plants in comparison to vector control plants during HS (Supplementary figure S9a, b). Fold change calculation of HSR genes by comparing their expression in TRV-EV and *TRV-HSFA7* silenced plants kept in HS revealed the down-regulation of these HSR genes (Supplementary figure S9a, b), indicating a positive role of *HSFA7a* in governing the transcription of HSR gene during HS.

**Figure 5:**
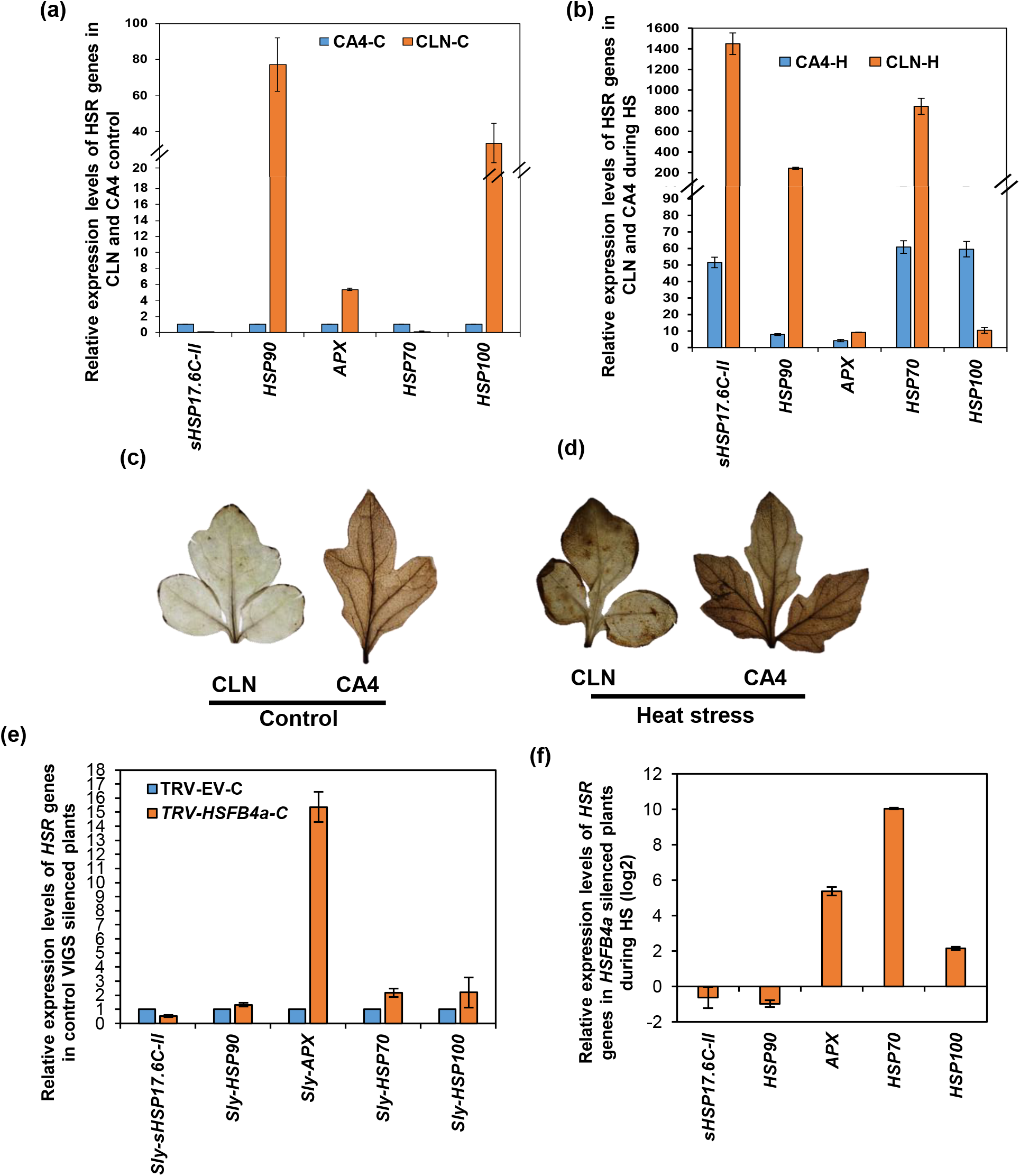
Expression patterns of heat stress responsive genes during heat stress in wild type and TRV-HSFB4a silenced plants. (a) Comparative expression profiles of HSR genes in CLN and CA4 tomato cultivars in control conditions. (b) Fold change induction of HSR genes in CLN and CA4 tomato cultivars during HS. Heat mediated induction of HSR genes is plotted as bar graph, the fold change is calculated by comparing the plants kept in control conditions and plants subjected to heat tress at 45°C. (c-d) DAB staining of control (c) and HS treated (d) CLN and CA4 plants to estimate the production of ROS. (e) Expression profiles of HSR genes in *TRV-EV* and *TRV-HSFB4a* silenced plants in control conditions. (f) qRT-PCR based fold change in expression levels of HSR genes in *TRV-EV* and *TRV-HSFB4a* silenced plants during heat stress. The expression levels of genes were calculated using the 2−ΔΔCt method and presented using fold-change values. The fold change normalization was done by setting the CA4-control value as one in (a), CA4 and CLN control as one in (b), TRV-EV-C as one in (e) and TRV-HS value as one in (f). Log2 transformation was applied to the fold-change data to obtain negative fold change values in (f). *Actin* was used as endogenous control. Error bars represent the standard error of independent biological replicates in (a, b, e and f).

Furthermore, to confirm the direct involvement of reduced *HSFB4a* levels in CLN in maintaining its higher HSR effectors levels and higher heat tolerance, we analysed the expression of these HSR genes in *TRV-HSFB4a* silenced plants. qRT-PCR based expression profiling of HSR genes in TRV-EV and *TRV-HSFB4a* silenced plants expression analysis revealed the higher expression of *HSP90, APX*, *HSP70* and *HSP100* genes in *TRV-HSFB4a* silenced plants in comparison to TRV-EV plants during control conditions (Figure 5e). The higher expression of HSR genes in *TRV-HSFB4a* silenced plants in control and heat stress conditions (Figure 5e, f) confirms *HSFB4a* as negative regulator of these HSR genes. Furthermore, to rule out the possibility that the HSR up-regulation in *TRV-HSFB4a* silenced plants during HS is a general heat stress response of the plants towards heat imposition and not due to *TRV-HSFB4a* silencing, we compared the expression of HSR genes in heat imposed TRV-EV and *TRV-HSFB4a* silenced plants. qRT-PCR based expression profiling revealed the strong up -regulation of *sHSP17.6C-II, APX*, *HSP70* and *HSP100* in *TRV-HSFB4a* silenced plants during heat stress (Supplementary figure S10), confirming the role of *HSFB4a* levels in governing the expression of HSR genes during HS. These findings support and validates the link between the reduced *HSFB4a* levels with high HSR gene expression in tolerant cultivar CLN. The higher HSR levels in CLN were maintained by the reduced expression of *HSFB4a.*

## Discussion

### Cultivar biased abundance of *HSFA7* and *HSFB4a* governs the heat stress tolerance of contrasting cultivars

Exploring the cultivar-biased regulation of genes is critical to unravel the key regulatory networks governing the tolerance mechanism under varied environmental conditions (Koeslin-Findekleeet al. 2014; Boccacci et al. 2017; Balyan et al. 2017). Several studies in literature report the differential regulation of gene expression in contrasting cultivar, where tolerant genotype have major extent of up-regulated events while sensitive cultivar show larger extent of down-regulated events (Frank et al. 2009; Balyan et al. 2020). Hu et al. (2020b) have shown that cultivated tomato germplasm has undergone a progressive loss of acclimation to strong temperature elevations. This sensitivity is associated with intronic polymorphisms in the HS transcription factor *HSFA2* which affect the splicing efficiency of its pre-mRNA. The authors show that intron splicing in wild species results in increased synthesis of an *HSFA2* isoform (*HSFA2-II*), implicated in the early stress response, at the expense of *HSFA2-I* which is involved in the establishment of short-term acclimation and thermotolerance. Recently, a study by Hu et al. (2020a) have shown that upon heat stress, the expression of majority of genes behaves similarly in all genotypes, including the HSFs and HSPs, genes involved in photosynthesis and mitochondrial ATP production. In turn, genes involved in hormone and RNA-based regulation, such as auxin- and ethylene-related genes and some transcription factors show a differential regulation that strongly associates with the cultivar thermotolerance pattern (Hu et al. 2020a). Another recent study by Balyan et al. (2020) has shown the orchestration of inherent cultivar-specific transcription factor cascade in regulating the transcription of key genes governing heat tolerance of tolerant tomato cultivar (CLN).

HSFs are the key transcription factors that regulate the organization of complex transcriptional cascade during HSR (Schramm et al. 2008; Yoshida et al. 2008, 2011; Chen et al. 2010; Sato et al. 2014). Therefore, to delineate the roles of HSFs in governing the tolerance of heat tolerant cultivar of tomato (CLN), we assessed the HS expression profiles of HSFs in heat tolerant (CLN) and sensitive (CA4) cultivars that are hot sub-tropical growing varieties. More than 55% of HSFs exhibit heat induced up-regulation in both cultivars, suggesting their common role in regulating HSR in both cultivars (Figure 1a, b). Several HSFs like *HSFA1b, HSFA2, HSFA3, HSFA7, HSFB1* and *HSFB2b* have also been annotated as central regulators that show enhanced transcript levels at 39 ◦C and 45 ◦C in all tolerant and sensitive genotypes of tomato (Hu et al. 2020a). In our study also, *HSFA7* exhibits very strong induction in CLN as compared to CA4 and we show *HSFA7* as a regulator of tomato heat stress response (Figure 2). Furthermore, *GUS* aided promoter reporter assays also confirmed the strong induction of *HSFA7* transcription during HS in the CLN background (Supplementary figure S2). Therefore, to explore the effector functions of *HSFA7* in governing heat tolerance of tomato, because of its strong induction during HS, we utilized the VIGS silencing and transient overexpression approach. Our analysis revealed that *HSFA7* acts as a positive regulator of thermotolerance, as plants with silenced *HSFA7* exhibit sensitivity to heat stress and a high degree of damage was scored for these silenced plants as compared to the vector control plants (Figure 2a, b). These results were further confirmed by transient over-expression of *HSFA7* in tomato seedlings, where P35S:*HSFA7* overexpressing plants exhibit high survival rate, enhanced hypocotyl length and higher biomass as compared to vector control plants in HS (Figure 2c-g). Larkindale and Vierling (2008) have also reported the role of *HSFA7* in regulating thermotolerance, as the T-DNA knockout Arabidopsis mutants of *HSFA7* had decreased thermotolerance as compared to wild-type plants. Balyan et al. (2020) have reported the inverse expression of *Notabilis*, a gene that regulates the rate limiting step in the ABA biosynthesis pathway in CLN and CA4 during HS. Silencing of *Notabilis* gene strongly reduces the transcription of *HSFA7* and *Notabilis silenced plants* exhibit heat sensitivity. These findings speculate that the differential level of ABA in CLN and CA4 contributed by inverse levels of *Notabilis* gene governs the differential levels of *HSFA7* induction in these cultivars. ABA-mediated regulation of *HSFA7* was further confirmed by the up-regulation of *HSFA7* in CLN in response to ABA application (Balyan et al. 2020). A link of HSFs and ABA response was also reported in *Arabidopsis thaliana,* where *HSFA6a* has been shown to regulate ABA related genes (Hwang et al. 2014). Furthermore, expression analysis of HSR genes in *TRV-HSFA7* silenced plants revealed the loss of heat-mediated induction of HSR genes during HS (Supplementary figure S9a, b). These findings established *HSFA7* as regulator of HS and involved in governing the thermotolerance of tomato.

Furthermore, we find that *HSFB4a* exhibits strong down-regulation in CLN (and not CA4) in response to heat stress (Figure 1a). A study by Ikeda and Ohme-Takagi (2009) has reported that class B HSFs contain an ‘R/KLFGV’ motif which acts as a transcriptional repressor. Therefore, this study further explored the cultivar biased transcriptional regulation of *HSFB4a* in CLN and CA4 backgrounds in HS by using *GUS* aided promoter reporter assays. This analysis revealed strong reduction of *GUS* transcripts during HS in CLN background and up-regulation in CA4 background, while the qRT-PCR expression data of *HSFB4a* in CA4 revealed that HSFB4 transcripts remain unchanged during HS (Figure 1a and Supplementary figure S6a, b). These results raised the possibility of additional regulation of *HSFB4a* transcripts. Indeed, we find a miRNA-mediated post-transcriptional regulation of *HSFB4a* by Sly-miR4200. *GFP* based transient *in-planta* assays confirm the Sly-miR4200-mediated cleavage of *HSFB4a* transcripts (Figure 3c, d). Expression profiling in CLN and CA4 during HS revealed the CLN specific strong up-regulation of *Sly-MIR4200*, while a slight up-regulation was observed in the CA4 background. Thus, we speculate that this differential abundance of *Sly-MIR4200* maintains the differential levels of *HSFB4a* between these cultivars in heat stress, thereby regulating the differential heat tolerance in CLN-CA4 cultivars (Supplementary figure S11). While microRNAs have been established as major regulators of transcription factors at post-transcriptional level in response to various abiotic stresses and developmental stages (Stief et al. 2014; Karlova et al. 2013; Chung et al. 2020; Rao et al. 2020), this is the first report to establish the miRNA-mediated post-transcriptional regulation of HSF by transcript cleavage. While the role of *Sly-miR4200:HSFB4a* is apparent in regulating thermotolerance mechanism in the tolerant and sensitive cultivars, further the expression analysis of *Sly-miR4200:HSFB4a* module also suggests the possible roles of this module in governing other abiotic stresses like drought and salt stress. Further functional studies using overexpression and knockdown approaches are required to assess their role in these abiotic stresses.

Furthermore, this study explored the functions of *HSFB4a* in regulating heat tolerance of CLN. VIGS and transient over-expression assays of *HSFB4a* confirm its negative role in governing heat tolerance (Figure 4). Also, expression analysis of HSR genes in *TRV-HSFB4a* silenced plants showed the strong up-regulation of *APX, HSP70* and *HSP100* genes during control and HS conditions (Figure 5e, f). This strong up-regulation of HSR genes could be correlated with the higher expression of these HSR genes in the tolerant cultivar CLN (Figure 5a, b), whereas the expression of *HSFB4a* is strongly reduced during control as well as HS condition (Figure 1a). Taking together our findings, the low transcripts levels of *HSFB4a* in tolerant cultivar CLN (Figure 1a) and in *TRV-HSFB4a* (Figure 4e) silenced plants could be correlated with the enhanced expression of *APX* gene, lower ROS production and higher survival of tolerant cultivar CLN (Figure 5a, b, c and d) and *TRV-HSFB4a* silenced plants (Figure 4i-j and figure 5e-f). These findings conclude that abundance of *HSFB4a* in CA4 may contribute towards its heat sensitivity by repressing the transcription of downstream genes like HSPs and APX. Hu et al. (2020a) have reported the expression of *HSFA6b* to relate with the tolerant/sensitive cultivar thermotolerance pattern in tomato. We on the other hand, do not find such a correlation for *HSFA6b* (Figure 1), instead show the role of *HSFB4a* in governing thermotolerance in a cultivar-specific manner. This indicates that tomato varieties growing in the temperate and sub-tropical climatic zones have evolved distinct thermotolerance regulatory mechanisms and different approaches may be needed to raise temperature-resilient plants based on their geographical location.

## Acknowledgments

NIPGR and DST-SERB, India (Project No.: EMR/2016/006229) funded and supported this work. The authors acknowledge phytotron facility, CIF and field area provided by NIPGR and AVRDC, Taiwan for providing CLN1621L and CA4 seeds. The authors are thankful to DBT-eLibrary Consortium (DeLCON) for providing access to e-resources. SR acknowledges Department of Biotechnology (DBT) Govt. of India and JRD acknowledges Council of Scientific and Industrial Research (CSIR) for the award of research fellowships.

## Author contributions

SM designed and supervised the experiments; SR designed and performed all the experiments and analysed the data; RV performed promoter reporter assays for promoter functionality. SR and SM wrote the article; JRD and SB helped in experimentation and figure preparation. SM agrees to serve as the author responsible for contact and ensures communication.

## Main Conclusion

Reduced *HSFB4a* repressor levels and enhanced *HSFA7* levels govern thermotolerance in the tomato. Cultivar-biased post-transcriptional regulation of *HSFB4a* by miR4200 mediates heat stress tolerance/sensitivity.

## Supporting material legends

**Supplementary figure S1: Relative abundance of *HSFA7* and *HSFB4a* transcripts in non-stressed leaf of contrasting tomato cultivars.**(a) Expression analysis of *HSFA7* and *HSFB4a* in leaf of CLN (tolerant) and CA4 (sensitive) cultivar by qRT-PCR. Fold change of *HSF* genes in control CLN plants is calculated by setting the fold change value of CA4 plants kept in control conditions as one. *Actin* was used as the normalization control. Experiment was repeated two times and average values are plotted as bars, error bars depict standard error between three replicates.

**Supplementary figure S2: Heat stress mediated regulation of *HSFA7a* transcription.**(a) *GUS* histochemical assay to assess transcriptional activity of *Pro-HSFA7:GUS* reporter construct during control and 2h of heat stress at 45°C in CLN. (b) Levels of *GUS* transcripts measured by qRT-PCR after agro-infiltration of *Pro-HSFA7:GUS* reporter constructs in tomato seedlings. All qRT-PCR experiments were repeated at least three times on 70 seedlings per replicate with similar results, and average data is shown. Error bar denotes standard error.

**Supplementary figure S3: Comparative promoter analysis of *HSFA7* between CLN and CA4.** Alignment of 2 kb up stream sequences from the ATG codon using CLC Genomics software.

**Supplementary figure S4: Virus induced silencing of tomato *HSFA7* and *PDS*. PDS used as silencing establishment control to analysis HSFA7 silencing.**(a) qRT-PCR analysis of *HSFA7* gene in TRV-EV and *TRV-HSFA7* plants confirming the silencing. The expression levels of genes were calculated using the 2−ΔΔCt method and presented using fold-change values. *Actin* was used for normalising expression. EV: Empty vector control. Error bars represent the standard error. (b) Silencing of the *PDS* control gene showing photobleaching in tomato leaves. Photographs were taken 3 weeks after infiltration of silencing constructs in (b).

**Supplementary figure S5: Comparative promoter analysis of *HSFB4a* between CLN and CA4.** Alignment of 2.5 kb up stream sequences from the ATG codon using CLC Genomics software.

**Supplementary figure S6: Heat stress mediated regulation of HSFB4a in leaf of contrasting cultivars.**(a-b) *GUS* expression patterns of pHSFB4a:*GUS* reporter constructs upon transient infiltration in CLN and CA4 background after exposing to 0 (control) or 2 h of heat stress at 45°C. For GUS reporter assay, 2 kb genomic sequences upstream of the translational start sites were used. Staining in (a) was performed 2 days after agro-infiltration in 4-days-old tomato seedlings post HS. All qRT-PCR experiments in (b) were repeated at least three times on 70 seedlings per replicate with similar results, and average data is shown. Error bar denotes standard error.

**Supplementary figure S7: Relative abundance of *Sly-MIR4200* in non-stressed leaf of contrasting tomato cultivars.**(a) Expression analysis of *Sly-MIR4200* in leaf of CLN (tolerant) and CA4 (sensitive) cultivar by qRT-PCR. Fold change of *Sly-MIR4200* in control CLN plants is calculated by setting the fold change value of CA4 plants kept in control conditions as one. *Actin* was used as the normalization control. Experiment was repeated two times and average values are plotted as bars, error bars depict standard error between three replicates.

**Supplementary figure S8: Determining the specificity of *TRV-HSFB4a* VIGS silencing in tomato plants.**(A) qRT-PCR based expression profiles of putative off target gene (HSFB4b) using the 2−ΔΔCt method in TRV-*HSFB4a* silenced plants. Error bars represent the standard error of biological replicates. *Actin* was used as endogenous control. The fold change normalization was done by setting the TRV-EV expression as one.

**Supplementary figure S9: Expression profiles of heat stress responsive genes during heat stress in *TRV-HSFA7* silenced plants.**(a) Comparative expression profiles of HSR genes in TRV-EV and *TRV-HSFA7* silenced plants during heat stress. Fold change of HSR genes in TRV-EV and *TRV-HSFA*7 silenced plants is calculated by setting the fold change value of TRV-EV and *TRV-HSFA*7 plants kept in control conditions as one. (b) Fold change induction of HSR genes in *TRV-HSFA*7 silenced tomato plants during HS. Heat mediated induction of HSR genes is plotted as bar graph, the fold change is calculated by comparing the TRV-EV and *TRV-HSFA*7 plants subjected to 4.5 hours of heat stress at 45°C. The expression levels of genes were calculated using the 2−ΔΔCt method and presented using fold-change values. The fold change normalization was done by setting the fold change value of TRV-EV-HS plants as one in (b). Log2 transformation was applied to the fold-change data to obtain negative fold change value in (b). *Actin* was used as endogenous control. Error bars represent the standard error of independent biological replicates in (a-b).

**Supplementary figure S10: Expression profiles of heat stress responsive genes during heat stress in *TRV-HSFB4a* silenced plants.**(a) Comparative expression profiles of HSR genes in TRV-EV and *TRV-HSFB4a* silenced plants during heat stress. Fold change of HSR genes in TRV-EV and *TRV-HSFB4a* silenced plants is calculated by setting the fold change value of TRV-EV and *TRV-HSFB4a* plants kept in control conditions as one. The expression levels of genes were calculated using the 2−ΔΔCt method and presented using fold-change values. *Actin* was used as endogenous control. Error bars represent the standard error of independent biological replicates.

**Supplementary figure S11: Transcriptional and miRNA mediated two tier regulation of *HSFB4a* in CLN and CA4.** Heat stress causes strong induction of Sly-miR4200 in CLN and very slight induction in CA4, this leads to differential accumulation *HSFB4a* transcript levels between CLN and CA4. Additional layer of regulation is executed at transcriptional level on *HSFB4a* by differential orchestration of cultivar specific TF modules in CLN and CA4. HSFB4a silencing leads to the enhancement of HSR genes like HSPs and ROS scavenger like APX resulting in less ROS burst and cell death in tolerant CLN cultivar. This cumulative regulation of *HSFB4a* in CLN and CA4 governs their thermotolerance. Red colored upward arrows depict the up-regulation of specific gene events, while green colored downward arrows depict down-regulation. Number of arrows depict the strength of up- and down regulation.

**Supplementary table S1:** Primers used in the study.

**Supplementary table S2:** Expression profiles of *HSFs* (log2 fold change values) during heat stress in contrasting cultivars.

**Supplementary table S3:** Summary of cis-regulatory elements present in HSFB4a promoter. For promoter analysis at PLANT-PAN version 3 tool, 2kb sequence upstream of the translational start site was used.

**Supplementary table S4:** Expression profiling of Sly-*MIR-4200* in CLN and CA4 during HS.

## References

Addo-Quaye, C., Miller, W. and Axtell, M.J., 2009. CleaveLand: a pipeline for using degradome data to find cleaved small RNA targets. Bioinformatics, 25(1), 130–131.

Balyan, S., Kumar, M., Mutum, R.D., Raghuvanshi, U., Agarwal, P., Mathur, S. and Raghuvanshi, S., 2017. Identification of miRNA-mediated drought responsive multitiered regulatory network in drought tolerant rice, Nagina 22. Scientific reports, 7(1), 1–17.

Balyan, S., Rao, S., Jha, S., Bansal, C., Das, J.R. and Mathur, S., 2020 Characterization of novel regulators for heat stress tolerance in tomato from Indian sub-continent. Plant Biotechnology Journal, doi:10.1111/pbi.13371.

Banti, V., Mafessoni, F., Loreti, E., Alpi, A. and Perata, P., 2010. The heat-inducible transcription factor HsfA2 enhances anoxia tolerance in Arabidopsis. Plant Physiology, 152(3), 1471–1483.

Bechtold, U., Albihlal, W.S., Lawson, T., Fryer, M.J., Sparrow, P.A., Richard, F., Persad, R., Bowden, L., Hickman, R., Martin, C. and Beynon, J.L., 2013. Arabidopsis HEAT SHOCK TRANSCRIPTION FACTORA1b overexpression enhances water productivity, resistance to drought, and infection. Journal of Experimental Botany, 64(11), 3467–3481.

Bita, C.E., Zenoni, S., Vriezen, W.H., Mariani, C., Pezzotti, M. and Gerats, T., 2011. Temperature stress differentially modulates transcription in meiotic anthers of heat-tolerant and heat-sensitive tomato plants. BMC genomics, 12(1), 384.

Boccacci, P., Mela, A., Mina, C.P., Chitarra, W., Perrone, I., Gribaudo, I. and Gambino, G., 2017. Cultivar-specific gene modulation in Vitisvinifera: analysis of the promoters regulating the expression of WOX transcription factors. Scientific reports, 7(1), 1–13.

Busch, W., Wunderlich, M. and Schöffl, F., 2005. Identification of novel heat shock factor-dependent genes and biochemical pathways in Arabidopsis thaliana. The Plant Journal, 41(1), 1–14.

Charng, Y.Y., Liu, H.C., Liu, N.Y., Chi, W.T., Wang, C.N., Chang, S.H. and Wang, T.T., 2007. A heat-inducible transcription factor, HsfA2, is required for extension of acquired thermotolerance in Arabidopsis. Plant physiology, 143(1), 251–262.

Charng, Y.Y., Liu, H.C., Liu, N.Y., Hsu, F.C. and Ko, S.S., 2006. Arabidopsis Hsa32, a novel heat shock protein, is essential for acquired thermotolerance during long recovery after acclimation. Plant Physiology, 140(4), 1297–1305.

Chen, H., Hwang, J.E., Lim, C.J., Kim, D.Y., Lee, S.Y. and Lim, C.O., 2010. Arabidopsis DREB2C functions as a transcriptional activator of HsfA3 during the heat stress response. Biochemical and biophysical research communications, 401(2), 238–244.

Chow, C.N., Lee, T.Y., Hung, Y.C., Li, G.Z., Tseng, K.C., Liu, Y.H., Kuo, P.L., Zheng, H.Q. and Chang, W.C., 2019. PlantPAN3. 0: a new and updated resource for reconstructing transcriptional regulatory networks from ChIP-seq experiments in plants. Nucleic acids research, 47(D1), D1155–D1163.

Chung, M.Y., Nath, U.K., Vrebalov, J., Gapper, N., Lee, J.M., Lee, D.J., Kim, C.K. and Giovannoni, J., 2020. Ectopic expression of miRNA172 in tomato (Solanum lycopersicum) reveals novel function in fruit development through regulation of an AP2 transcription factor. BMC plant biology, 20(1), 1–15.

Cortijo, S., Charoensawan, V., Brestovitsky, A., Buning, R., Ravarani, C., Rhodes, D., van Noort, J., Jaeger, K.E. and Wigge, P.A., 2017. Transcriptional regulation of the ambient temperature response by H2A. Z nucleosomes and HSF1 transcription factors in Arabidopsis. Molecular plant, 10(10), 1258–1273.

Daudi, A., Cheng, Z., O’Brien, J.A., Mammarella, N., Khan, S., Ausubel, F.M. and Bolwell, G.P., 2012. The apoplastic oxidative burst peroxidase in Arabidopsis is a major component of pattern-triggered immunity. The Plant Cell, 24(1), 275–287.

Fragkostefanakis, S., Mesihovic, A., Simm, S., Paupière, M.J., Hu, Y., Paul, P., Mishra, S.K., Tschiersch, B., Theres, K., Bovy, A. and Schleiff, E., 2016. HsfA2 controls the activity of developmentally and stress-regulated heat stress protection mechanisms in tomato male reproductive tissues. Plant physiology, 170(4), 2461–2477.

Frank, G., Pressman, E., Ophir, R., Althan, L., Shaked, R., Freedman, M., Shen, S. and Firon, N., 2009. Transcriptional profiling of maturing tomato (Solanum lycopersicum L.) microspores reveals the involvement of heat shock proteins, ROS scavengers, hormones, and sugars in the heat stress response. Journal of experimental botany, 60(13), 3891–3908.

Giorno, F., Wolters-Arts, M., Grillo, S., Scharf, K.D., Vriezen, W.H. and Mariani, C., 2010. Developmental and heat stress-regulated expression of HsfA2 and small heat shock proteins in tomato anthers. Journal of experimental botany, 61(2), 453–462.

Han, N., Fan, S., Zhang, T., Sun, H., Zhu, Y., Gong, H. and Guo, J., 2020. SlHY5 is a necessary regulator of the cold acclimation response in tomato. Plant Growth Regulation, 1–12.

Hu, Y., Fragkostefanakis, S., Schleiff, E. and Simm, S., 2020a. Transcriptional Basis for Differential Thermosensitivity of Seedlings of Various Tomato Genotypes. Genes, 11(6), 655.

Hu, Y., Mesihovic, A., Jiménez-Gómez, J.M., Röth, S., Gebhardt, P., Bublak, D., Bovy, A., Scharf, K.D., Schleiff, E. and Fragkostefanakis, S., 2020b. Natural variation in HsfA2 pre-mRNA splicing is associated with changes in thermotolerance during tomato domestication. New Phytologist, 225(3), 1297–1310.

Huang, Y.C., Niu, C.Y., Yang, C.R. and Jinn, T.L., 2016. The heat stress factor HSFA6b connects ABA signaling and ABA-mediated heat responses. Plant physiology, 172(2),1182–1199.

Hwang, S.M., Kim, D.W., Woo, M.S., Jeong, H.S., Son, Y.S., Akhter, S., Choi, G.J. and Bahk, J.D., 2014. Functional characterization of A rabidopsis HsfA6a as a heat-shock transcription factor under high salinity and dehydration conditions. Plant, cell & environment, 37(5), 1202–1222.

Ikeda, M. and Ohme-Takagi, M., 2009. A novel group of transcriptional repressors in Arabidopsis. Plant and cell physiology, 50(5), 970–975.

Ikeda, M., Mitsuda, N. and Ohme-Takagi, M., 2011. Arabidopsis HsfB1 and HsfB2b act as repressors of the expression of heat-inducible Hsfs but positively regulate the acquired thermotolerance. Plant physiology, 157(3), 1243–1254.

Karlova, R., van Haarst, J.C., Maliepaard, C., van de Geest, H., Bovy, A.G., Lammers, M., Angenent, G.C. and de Maagd, R.A., 2013. Identification of microRNA targets in tomato fruit development using high-throughput sequencing and degradome analysis. Journal of experimental botany, 64(7), 1863–1878.

Koch, E. and Slusarenko, A., 1990. Arabidopsis is susceptible to infection by a downy mildew fungus. The Plant Cell, 2(5), 437–445.

Koeslin-Findeklee, F., Meyer, A., Girke, A., Beckmann, K. and Horst, W.J., 2014. The superior nitrogen efficiency of winter oilseed rape (Brassica napus L.) hybrids is not related to delayed nitrogen starvation-induced leaf senescence. Plant and Soil, 384(1-2), 347–362.

Kotak, S., Larkindale, J., Lee, U., von Koskull-Döring, P., Vierling, E. and Scharf, K.D., 2007. Complexity of the heat stress response in plants. Current opinion in plant biology, 10(3), 310–316.

Kumar, M., Busch, W., Birke, H., Kemmerling, B., Nürnberger, T., Schöffl, F., 2009. Heat shock factors HsfB1 and HsfB2b are involved in the regulation of Pdf1.2 expression and pathogen resistance in Arabidopsis. Mol Plant., 2(1):152–165.

Larkindale, J. and Vierling, E., 2008. Core genome responses involved in acclimation to high temperature. Plant physiology, 146(2), 748–761.

Li, Z., Palmer, W.M., Martin, A.P., Wang, R., Rainsford, F., Jin, Y., Patrick, J.W., Yang, Y. and Ruan, Y.L., 2012. High invertase activity in tomato reproductive organs correlates with enhanced sucrose import into, and heat tolerance of, young fruit. Journal of experimental botany, 63(3), 1155–1166.

Liu, A.L., Zou, J., Liu, C.F., Zhou, X.Y., Zhang, X.W., Luo, G.Y. and Chen, X.B., 2013. Over-expression of OsHsfA7 enhanced salt and drought tolerance in transgenic rice. BMB reports, 46(1), 31.

Liu, H.C. and Charng, Y.Y., 2013. Common and distinct functions of Arabidopsis class A1 and A2 heat shock factors in diverse abiotic stress responses and development. Plant physiology, 163(1), 276–290.

Liu, H.C., Liao, H.T. and Charng, Y.Y., 2011. The role of class A1 heat shock factors (HSFA1s) in response to heat and other stresses in Arabidopsis. Plant, cell & environment, 34(5), 738–751.

Mishra, S.K., Tripp, J., Winkelhaus, S., Tschiersch, B., Theres, K., Nover, L. and Scharf, K.D., 2002. In the complex family of heat stress transcription factors, HsfA1 has a unique role as master regulator of thermotolerance in tomato. Genes & Development, 16(12), 1555–1567.

Mittler, R., 2006. Abiotic stress, the field environment and stress combination. Trends in plant science, 11(1), 15–19.

Nishizawa, A., Yabuta, Y., Yoshida, E., Maruta, T., Yoshimura, K. and Shigeoka, S., 2006. Arabidopsis heat shock transcription factor A2 as a key regulator in response to several types of environmental stress. The Plant Journal, 48(4), 535–547.

Nishizawa-Yokoi, A., Nosaka, R., Hayashi, H., Tainaka, H., Maruta, T., Tamoi, M., Ikeda, M., Ohme-Takagi, M., Yoshimura, K., Yabuta, Y. and Shigeoka, S., 2011. HsfA1d and HsfA1e involved in the transcriptional regulation of HsfA2 function as key regulators for the Hsfsignaling network in response to environmental stress. Plant and Cell Physiology, 52(5), 933–945.

Ohama, N., Kusakabe, K., Mizoi, J., Zhao, H., Kidokoro, S., Koizumi, S., Takahashi, F., Ishida, T., Yanagisawa, S., Shinozaki, K. and Yamaguchi-Shinozaki, K., 2016. The transcriptional cascade in the heat stress response of Arabidopsis is strictly regulated at the level of transcription factor expression. The Plant Cell, 28(1), 181–201.

Paul, A., Rao, S. and Mathur, S., 2016. The α-crystallin domain containing genes: identification, phylogeny and expression profiling in abiotic stress, phytohormone response and development in tomato (Solanum lycopersicum). Frontiers in plant science, 7, 426.

Qiu, Z., Wang, X., Gao, J., Guo, Y., Huang, Z. and Du, Y., 2016. The tomato Hoffman’s anthocyaninless gene encodes a bHLH transcription factor involved in anthocyanin biosynthesis that is developmentally regulated and induced by low temperatures. PloS one, 11, e0151067

Queitsch, C., Hong, S.W., Vierling, E. and Lindquist, S., 2000. Heat shock protein 101 plays a crucial role in thermotolerance in Arabidopsis. The Plant Cell, 12(4), 479–492.

Rao, S., Balyan, S., Jha, S. and Mathur, S., 2020. Novel insights into expansion and functional diversification of MIR169 family in tomato. Planta, 251(2), 55.

Rao, S., Bansal, C., Sorin, C., Crespi, M., Mathur, S., 2021. A conserved HSF:miR169:NF-YA loop involved in tomato and Arabidopsis heat stress tolerance. doi: https://doi.org/10.1101/2021.01.01.425064.

Rombauts, S., Déhais, P., Van Montagu, M. and Rouzé, P., 1999. PlantCARE, a plant cis-acting regulatory element database. Nucleic acids research, 27(1), 295–296.

Rivero, R.M., Ruiz, J.M., Garcıa, P.C., Lopez-Lefebre, L.R., Sánchez, E. and Romero, L., 2001. Resistance to cold and heat stress: accumulation of phenolic compounds in tomato and watermelon plants. Plant Science, 160(2), 315–321.

Sakuma, Y., Maruyama, K., Qin, F., Osakabe, Y., Shinozaki, K. and Yamaguchi-Shinozaki, K., 2006. Dual function of an Arabidopsis transcription factor DREB2A in water-stress-responsive and heat-stress-responsive gene expression. Proceedings of the National Academy of Sciences, 103(49), 18822–18827.

Sato, H., Mizoi, J., Tanaka, H., Maruyama, K., Qin, F., Osakabe, Y., Morimoto, K., Ohori, T., Kusakabe, K., Nagata, M. and Shinozaki, K., 2014. Arabidopsis DPB3-1, a DREB2A interactor, specifically enhances heat stress-induced gene expression by forming a heat stress-specific transcriptional complex with NF-Y subunits. The Plant Cell, 26(12), 4954–4973.

Sato, S., Kamiyama, M., Iwata, T., Makita, N., Furukawa, H. and Ikeda, H., 2006. Moderate increase of mean daily temperature adversely affects fruit set of Lycopersiconesculentum by disrupting specific physiological processes in male reproductive development. Annals of Botany, 97(5), 731–738.

Scharf, K.D., Berberich, T., Ebersberger, I. and Nover, L., 2012. The plant heat stress transcription factor (Hsf) family: structure, function and evolution. Biochimica et BiophysicaActa (BBA)-Gene Regulatory Mechanisms, 1819(2), 104–119.

Schramm, F., Ganguli, A., Kiehlmann, E., Englich, G., Walch, D. and von Koskull-Döring, P., 2006. The heat stress transcription factor HsfA2 serves as a regulatory amplifier of a subset of genes in the heat stress response in Arabidopsis. Plant molecular biology, 60(5), 759–772.

Schramm, F., Larkindale, J., Kiehlmann, E., Ganguli, A., Englich, G., Vierling, E. and von Koskull-Döring, P., 2008. A cascade of transcription factor DREB2A and heat stress transcription factor HsfA3 regulates the heat stress response of Arabidopsis. The Plant Journal, 53(2), 264–274.

Senthil-Kumar, M. and Mysore, K.S., 2014. Tobacco rattle virus–based virus-induced gene silencing in Nicotiana benthamiana. Nature protocols, 9(7), 1549–1562.

Stief, A., Altmann, S., Hoffmann, K., Pant, B.D., Scheible, W.R. and Bäurle, I., 2014. Arabidopsis miR156 regulates tolerance to recurring environmental stress through SPL transcription factors. The Plant Cell, 26(4), 1792–1807.

Sugio, A., Dreos, R., Aparicio, F. and Maule, A.J., 2009. The cytosolic protein response as a subcomponent of the wider heat shock response in Arabidopsis. The Plant Cell, 21(2), 642–654.

Suzuki, N., Miller, G., Morales, J., Shulaev, V., Torres, M.A. and Mittler, R., 2011. Respiratory burst oxidases: the engines of ROS signaling. Current opinion in plant biology, 14(6), 691–699.

Tang, M., Xu, L., Wang, Y., Cheng, W., Luo, X., Xie, Y., Fan, L. and Liu, L., 2019. Genome-wide characterization and evolutionary analysis of heat shock transcription factors (HSFs) to reveal their potential role under abiotic stresses in radish (Raphanussativus L.). BMC genomics, 20(1),1–13.

von Koskull-Döring, P., Scharf, K.D. and Nover, L., 2007. The diversity of plant heat stress transcription factors. Trends in plant science, 12(10), 452–457.

Waleed, S.A., Obomighie, I., Blein, T., Persad, R., Chernukhin, I., Crespi, M., Bechtold, U. and Mullineaux, P.M., 2018. Arabidopsis HEAT SHOCK TRANSCRIPTION FACTORA1b regulates multiple developmental genes under benign and stress conditions. Journal of Experimental Botany, 69(11), 2847–2862.

Wei, Y., Liu, G., Chang, Y., He, C. and Shi, H., 2018. Heat shock transcription factor 3 regulates plant immune response through modulation of salicylic acid accumulation and signalling in cassava. Molecular plant pathology, 19(10), 2209–2220.

Wunderlich, M., Doll, J., Busch, W., Kleindt, C.K., Lohmann, C. and Schöffl, F., 2007. Heat shock factors: regulators of early and late functions in plant stress response. Plant Stress, 1(1), 16–22.

Xin, M., Peng, H., Ni, Z., Yao, Y., Hu, Z. and Sun, Q., 2019. Wheat responses and tolerance to high temperature. In wheat production in changing environments (139–147). Springer, Singapore.

Yan, M.Y., Xie, D.L., Cao, J.J., Xia, X.J., Shi, K., Zhou, Y.H., Zhou, J., Foyer, C.H. and Yu, J.Q., 2020. Brassinosteroid-mediated reactive oxygen species are essential for tapetum degradation and pollen fertility in tomato. The Plant Journal, 102(5), 931–947.

Yoshida, T., Ohama, N., Nakajima, J., Kidokoro, S., Mizoi, J., Nakashima, K., Maruyama, K., Kim, J.M., Seki, M., Todaka, D. and Osakabe, Y., 2011. Arabidopsis HsfA1 transcription factors function as the main positive regulators in heat shock-responsive gene expression. Molecular Genetics and Genomics, 286(5-6), 321–332.

Yoshida, T., Sakuma, Y., Todaka, D., Maruyama, K., Qin, F., Mizoi, J., Kidokoro, S., Fujita, Y., Shinozaki, K. and Yamaguchi-Shinozaki, K., 2008. Functional analysis of an Arabidopsis heat-shock transcription factor HsfA3 in the transcriptional cascade downstream of the DREB2A stress-regulatory system. Biochemical and biophysical research communications, 368(3), 515–521.

Zang, D., Wang, J., Zhang, X., Liu, Z. and Wang, Y., 2019. Arabidopsis heat shock transcription factor HSFA7b positively mediates salt stress tolerance by binding to an E-box-like motif to regulate gene expression. Journal of experimental botany, 70(19), 5355–5374.

