## Supplementary figures for "Transcriptional regulation of *HSFA7* and post-transcriptional modulation of *HSFB4a* by miRNA4200 govern general and varietal thermotolerance in tomato"

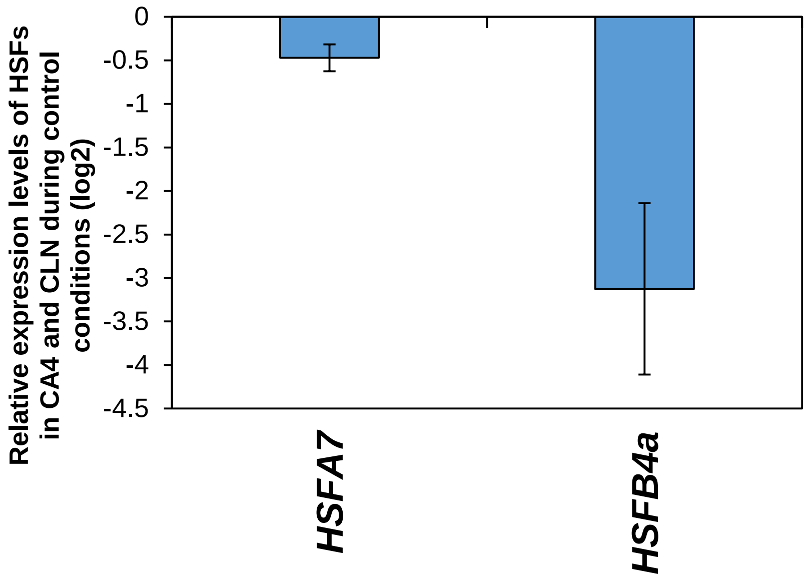


**Supplementary figure S1: Relative abundance of *HSFA7* and *HSFB4a* transcripts in non-stressed leaf of contrasting tomato cultivars.** (a) Expression analysis of *HSFA7* and *HSFB4a* in leaf of CLN (tolerant) and CA4 (sensitive) cultivar by qRT-PCR. Fold change of *HSF* genes in control CLN plants is calculated by setting the fold change value of CA4 plants kept in control conditions as one. *Actin* was used as the normalization control. Experiment was repeated two times and average values are plotted as bars, error bars depict standard error between three replicates.

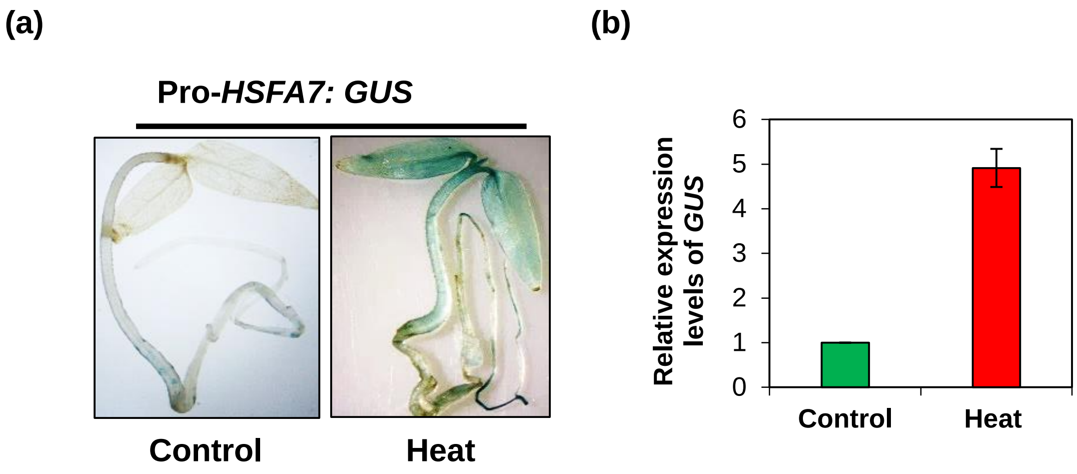


**Supplementary figure S2: Heat stress mediated regulation of *HSFA7a* transcription.** (a) *GUS* histochemical assay to assess transcriptional activity of *Pro-HSFA7:GUS* reporter construct during control and 2h of heat stress at 45°C in CLN. (b) Levels of *GUS* transcripts measured by qRT-PCR after agro-infiltration of *Pro-HSFA7:GUS* reporter constructs in tomato seedlings. All qRT-PCR experiments were repeated at least three times on 70 seedlings per replicate with similar results, and average data is shown. Error bar denotes standard error.

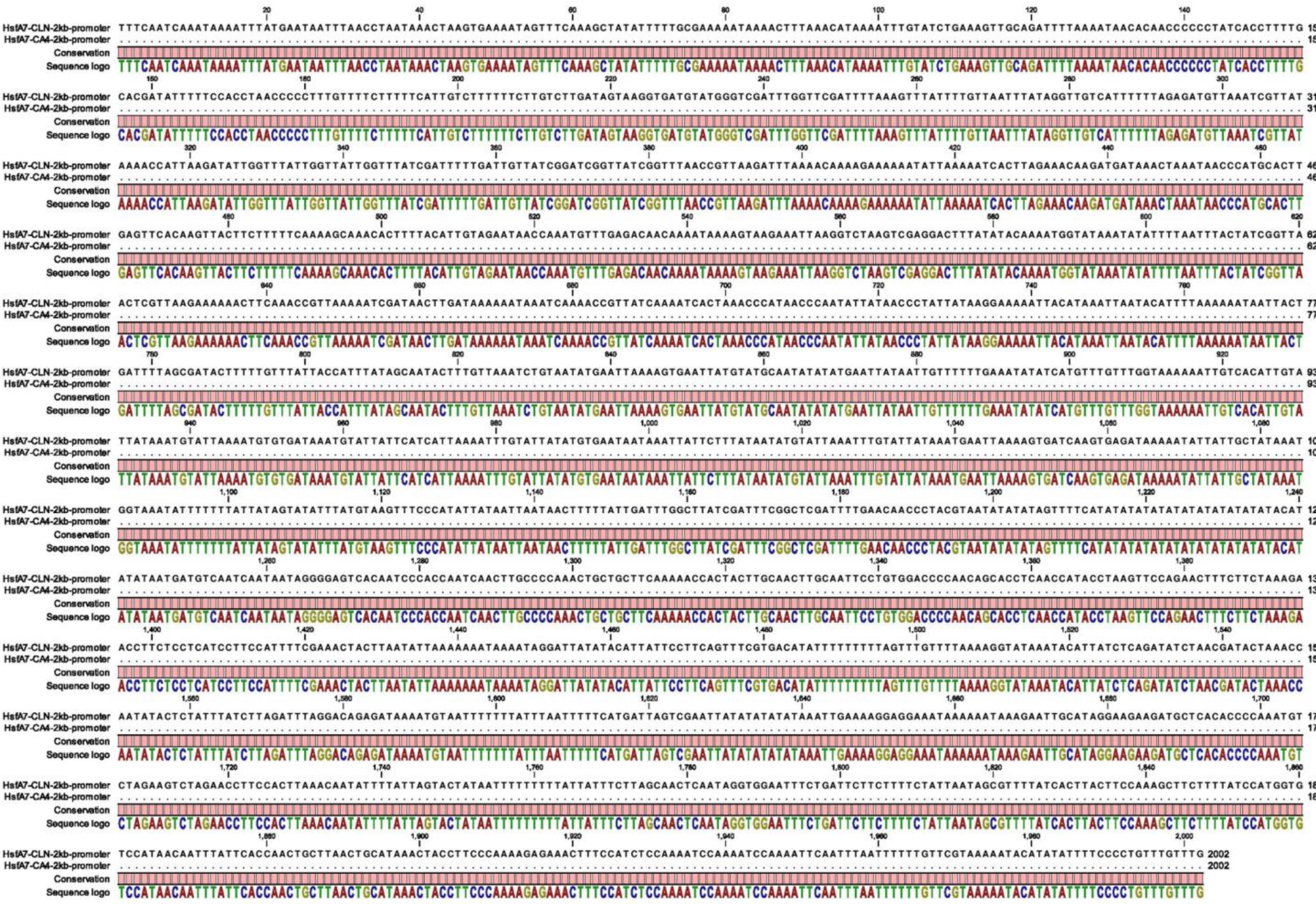

**Supplementary figure S3: Comparative promoter analysis of *HSFA7* between CLN and CA4.** Alignment of 2 kb up stream sequences from the ATG codon using CLC Genomics software.

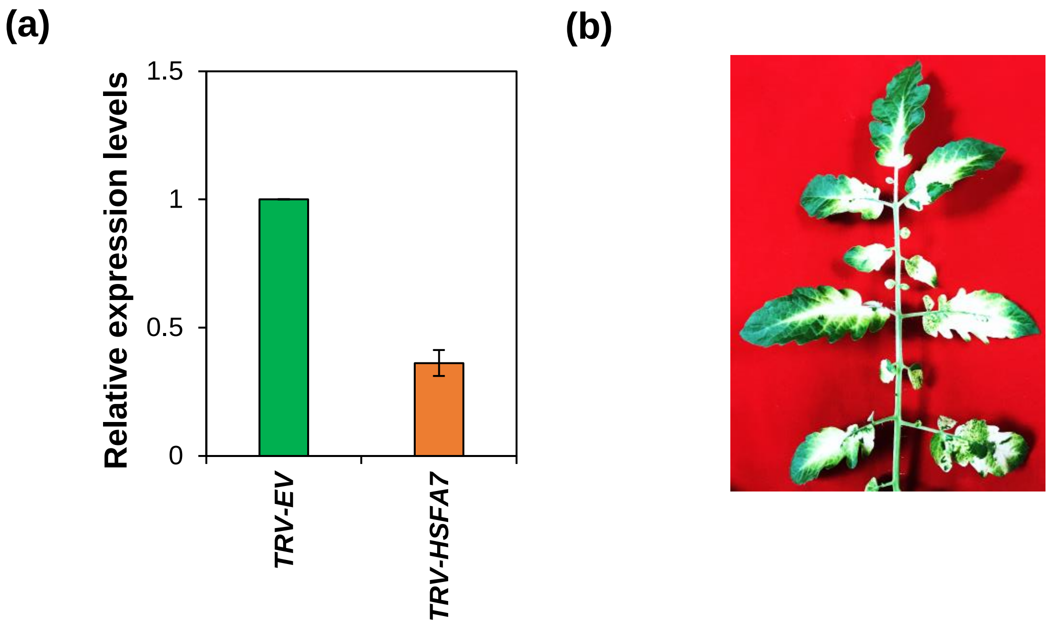


**Supplementary figure S4: Virus induced silencing of tomato *HSFA7* and *PDS*.** PDS used as silencing establishment control to analysis HSFA7 silencing. (a) qRT-PCR analysis of *HSFA7* gene in TRV-EV and *TRV-HSFA7* plants confirming the silencing. The expression levels of genes were calculated using the $2^{-\Delta\Delta Ct}$ method and presented using fold-change values. *Actin* was used for normalising expression. EV: Empty vector control. Error bars represent the standard error. (b) Silencing of the *PDS* control gene showing photobleaching in tomato leaves. Photographs were taken 3 weeks after infiltration of silencing constructs in (b).

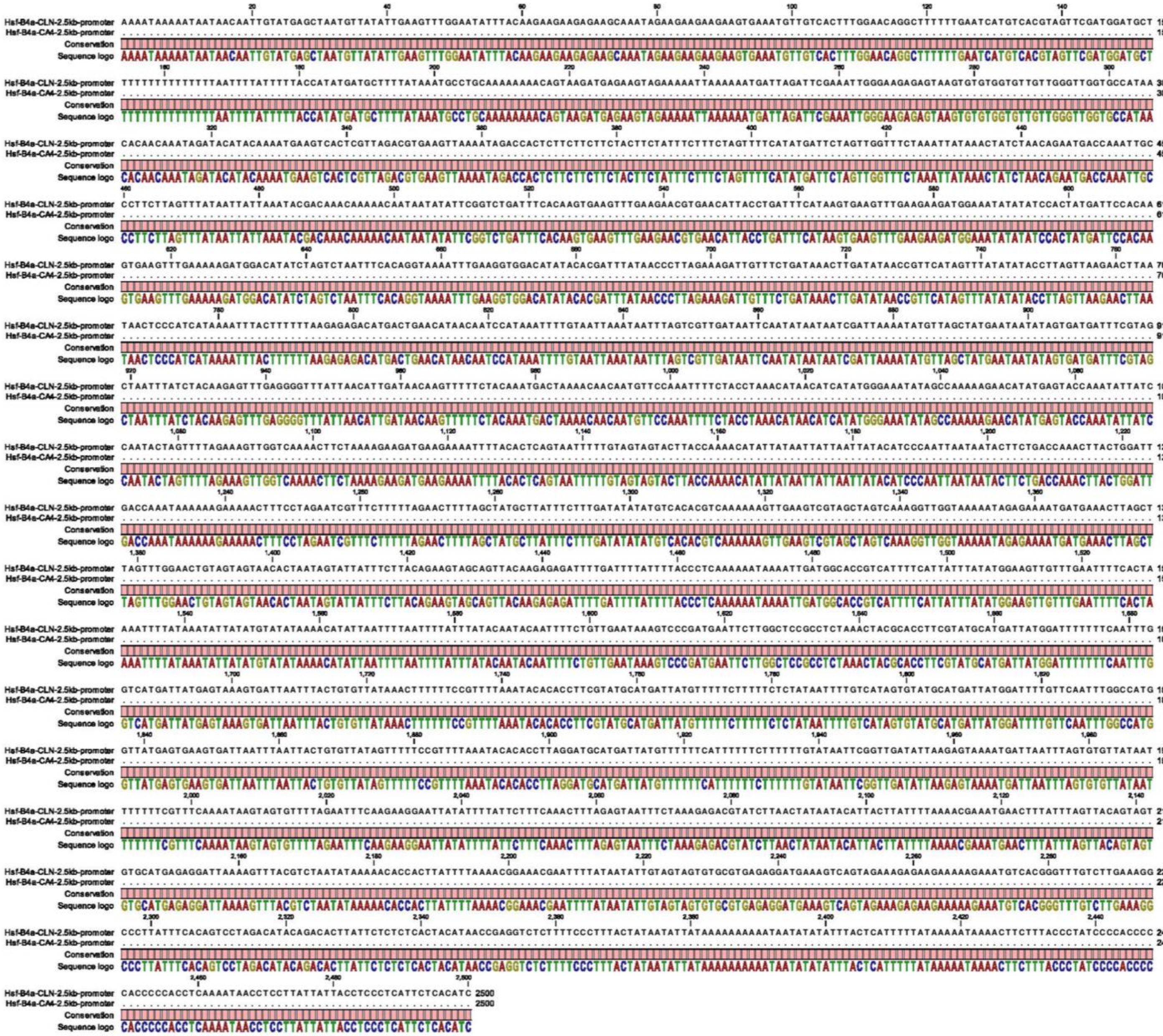

**Supplementary figure S5: Comparative promoter analysis of *HSFB4a* between CLN and CA4.** Alignment of 2.5 kb up stream sequences from the ATG codon using CLC Genomics software.

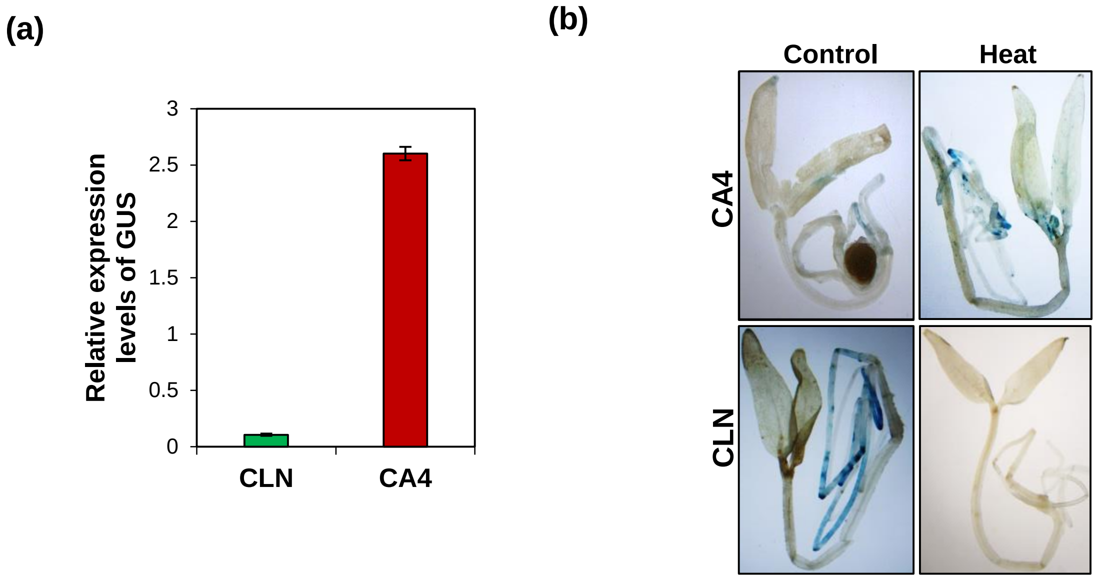


**Supplementary figure S6: Heat stress mediated regulation of HSFB4a in leaf of contrasting cultivars.** (a-b**)** *GUS* expression patterns of pHSFB4a:*GUS* reporter constructs upon transient infiltration in CLN and CA4 background after exposing to 0 (control) or 2 h of heat stress at 45°C. For GUS reporter assay, 2 kb genomic sequences upstream of the translational start sites were used. Staining in (a) was performed 2 days after agro-infiltration in 4-days-old tomato seedlings post HS. All qRT-PCR experiments in (b) were repeated at least three times on 70 seedlings per replicate with similar results, and average data is shown. Error bar denotes standard error.

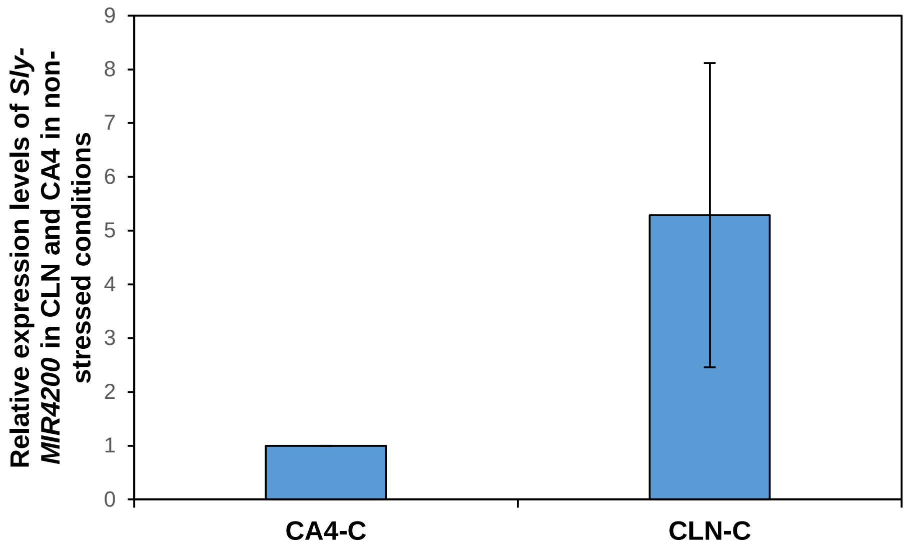


**Supplementary figure S7: Relative abundance of *Sly-MIR4200* in non-stressed leaf of contrasting tomato cultivars.** (a) Expression analysis of *Sly-MIR4200* in leaf of CLN (tolerant) and CA4 (sensitive) cultivar by qRT-PCR. Fold change of *Sly-MIR4200* in control CLN plants is calculated by setting the fold change value of CA4 plants kept in control conditions as one. *Actin* was used as the normalization control. Experiment was repeated two times and average values are plotted as bars, error bars depict standard error between three replicates.

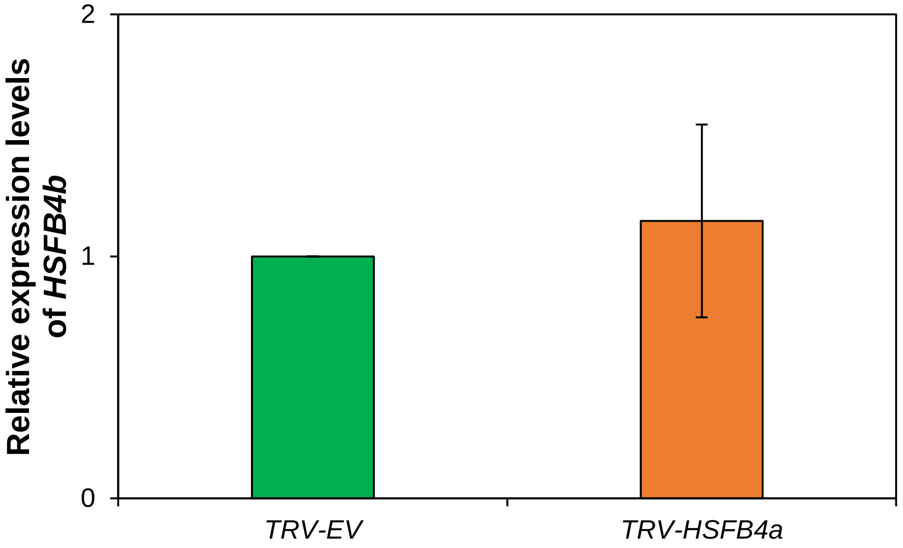


**Supplementary figure S8: Determining the specificity of *TRV-HSFB4a* VIGS silencing in tomato plants.** (A) qRT-PCR based expression profiles of putative off target gene (HSFB4b) using the $2^{-\Delta\Delta Ct}$ method in TRV-*HSFB4a* silenced plants. Error bars represent the standard error of biological replicates. *Actin* was used as endogenous control. The fold change normalization was done by setting the TRV-EV expression as one.

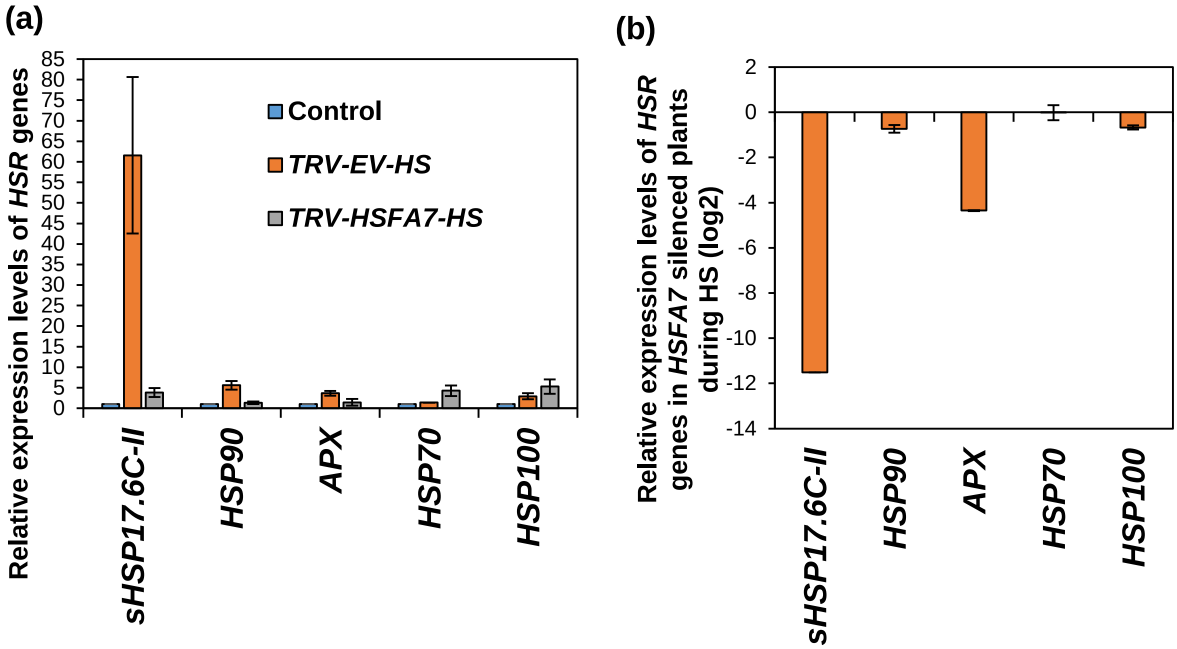


**Supplementary figure S9 : Expression profiles of heat stress responsive genes during heat stress in *TRV-HSFA7* silenced plants.** (a) Comparative expression profiles of HSR genes in TRV-EV and *TRV-HSFA7* silenced plants during heat stress. Fold change of HSR genes in TRV-EV and *TRV-HSFA*7 silenced plants is calculated by setting the fold change value of TRV-EV and *TRV-HSFA*7 plants kept in control conditions as one. (b) Fold change induction of HSR genes in *TRV-HSFA*7 silenced tomato plants during HS. Heat mediated induction of HSR genes is plotted as bar graph, the fold change is calculated by comparing the TRV-EV and *TRV-HSFA*7 plants subjected to 4.5 hours of heat stress at 45°C. The expression levels of genes were calculated using the $2^{-\Delta\Delta Ct}$ method and presented using fold-change values. The fold change normalization was done by setting the fold change value of TRV-EV-HS plants as one in (b). Log2 transformation was applied to the fold-change data to obtain negative fold change value in (b). *Actin* was used as endogenous control. Error bars represent the standard error of independent biological replicates in (a-b).

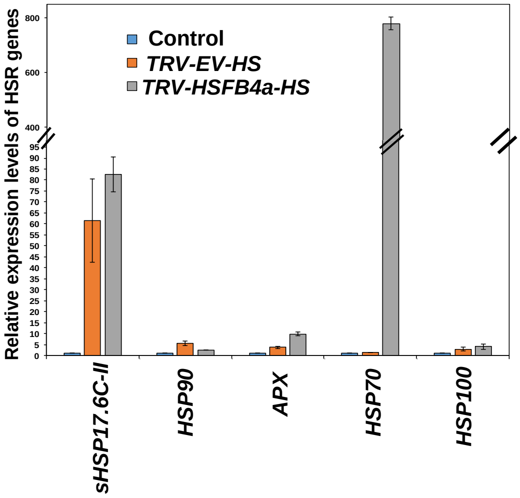


**Supplementary figure S10 : Expression profiles of heat stress responsive genes during heat stress in *TRV-HSFB4a* silenced plants.** (a) Comparative expression profiles of HSR genes in TRV-EV and *TRV-HSFB4a* silenced plants during heat stress. Fold change of HSR genes in TRV-EV and *TRV-HSFB4a* silenced plants is calculated by setting the fold change value of TRV-EV and *TRV-HSFB4a* plants kept in control conditions as one. The expression levels of genes were calculated using the $2^{-\Delta\Delta Ct}$ method and presented using fold-change values. *Actin* was used as endogenous control. Error bars represent the standard error of independent biological replicates.

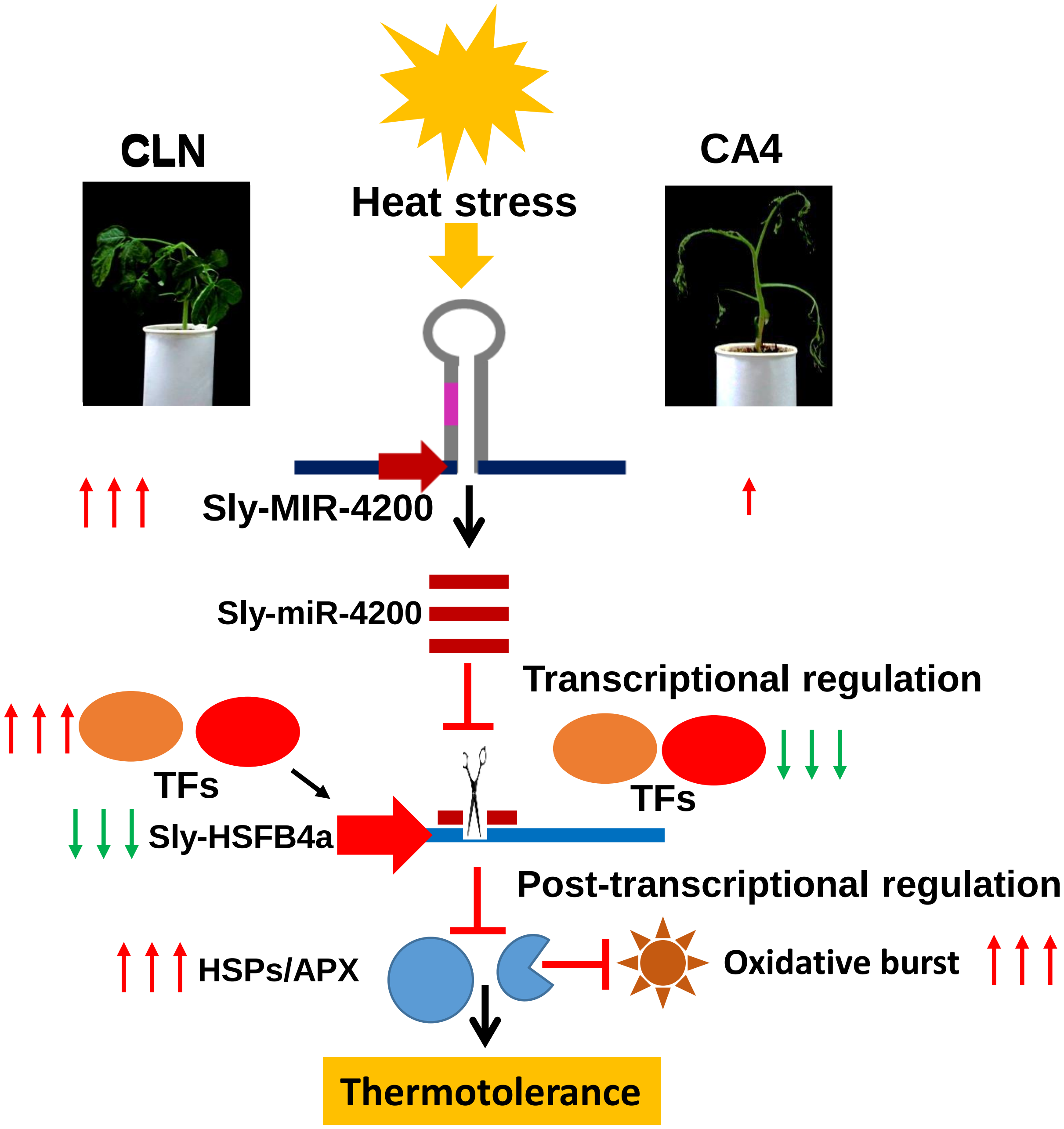


**Supplementary figure S11: Transcriptional and miRNA mediated two tier regulation of *HSFB4a* in CLN and CA4.** Heat stress causes strong induction of Sly-miR4200 in CLN and very slight induction in CA4, this leads to differential accumulation *HSFB4a* transcript levels between CLN and CA4. Additional layer of regulation is executed at transcriptional level on *HSFB4a* by differential orchestration of cultivar specific TF modules in CLN and CA4. HSFB4a silencing leads to the enhancement of HSR genes like HSPs and ROS scavenger like APX resulting in less ROS burst and cell death in tolerant CLN cultivar. This cumulative regulation of *HSFB4a* in CLN and CA4 governs their thermotolerance. Red colored upward arrows depict the up-regulation of specific gene events, while green colored downward arrows depict down-regulation. Number of arrows depict the strength of up- and down regulation.
